# Electrophysiological correlates of perceptual prediction error are attenuated in dyslexia

**DOI:** 10.1101/2021.06.22.449408

**Authors:** Sara D. Beach, Sung-Joo Lim, Carlos Cardenas-Iniguez, Marianna D. Eddy, John D. E. Gabrieli, Tyler K. Perrachione

**Affiliations:** McGovern Institute for Brain Research, Massachusetts Institute of Technology 77 Massachusetts Avenue, Cambridge, MA 02139 U.S.A.; Department of Brain and Cognitive Sciences, Massachusetts Institute of Technology 77 Massachusetts Avenue, Cambridge, MA 02139 U.S.A.; Program in Speech and Hearing Bioscience and Technology, Harvard University 260 Longwood Avenue, Boston, MA 02115 U.S.A.; Department of Speech, Language, and Hearing Sciences, Boston University 635 Commonwealth Avenue, Boston, MA 02215 U.S.A.

**Keywords:** prediction error, repetition, expectation, adaptation, dyslexia, event-related potentials, time-frequency

## Abstract

A perceptual adaptation deficit often accompanies reading difficulty in dyslexia, manifesting in poor perceptual learning of consistent stimuli and reduced neurophysiological adaptation to stimulus repetition. However, it is not known how adaptation deficits relate to differences in feedforward or feedback processes in the brain. Here we used electroencephalography (EEG) to interrogate the feedforward and feedback contributions to neural adaptation as adults with and without dyslexia viewed pairs of faces and words in a paradigm that manipulated whether there was a high probability of stimulus repetition versus a high probability of stimulus change. We measured three neural dependent variables: *expectation* (the difference between prestimulus EEG power with and without the expectation of stimulus repetition), feedforward *repetition* (the difference between event-related potentials (ERPs) evoked by an expected change and an unexpected repetition), and feedback-mediated *prediction error* (the difference between ERPs evoked by an unexpected change and an expected repetition). Expectation significantly modulated prestimulus theta- and alpha-band EEG in both groups. Unexpected repetitions of words, but not faces, also led to significant feedforward repetition effects in the ERPs of both groups. However, neural prediction error when an unexpected change occurred instead of an expected repetition was significantly weaker in dyslexia than the control group for both faces and words. These results suggest that the neural and perceptual adaptation deficits observed in dyslexia reflect the failure to effectively integrate perceptual predictions with feedforward sensory processing. In addition to reducing perceptual efficiency, the attenuation of neural prediction error signals would also be deleterious to the wide range of perceptual and procedural learning abilities that are critical for developing accurate and fluent reading skills.

## 1. Introduction

Dyslexia is a developmental disorder characterized by poor reading skills despite adequate educational opportunity (Lyon, Shaywitz, & Shaywitz, 2003). Numerous studies have documented structural and functional alterations in the brains of individuals with dyslexia (reviewed in Gabrieli, 2009; Linkersdörfer et al., 2012; Martin, Kronbichler, & Richlan, 2016). Fluent reading involves the integration of visual and linguistic processes, is supported by attention and memory, and may overtly or covertly engage auditory and motor systems. This masterful orchestration has appeared too recently in human culture for it to be shaped by the pressures of natural selection, and therefore the brain’s reading network has been described as one that recycles circuits that subserve evolutionarily older functions (Dehaene & Cohen, 2007). The challenges to developing fluent reading skills in dyslexia must therefore come from latent dysfunction in either the circuits that develop into the reading network or the plasticity processes that support repurposing those circuits for reading. In this study, we investigated the neural bases of differences in rapid perceptual adaptation in dyslexia—a recently documented, domain-general deficit that may reflect weakness in the plasticity processes that support learning to read (Ahissar et al., 2006; Gabay & Holt, 2020; Jaffe-Dax et al., 2016; Oganian & Ahissar, 2012). Specifically, we aimed to expand on the evidence for diminished neural adaptation in dyslexia (Perrachione et al., 2016) by determining whether this phenomenon is due to differences in bottom-up feedforward processing, top-down expectations, or their interaction.

### 1.1 Neural and behavioral adaptation deficits in dyslexia

Neural systems take advantage of consistent sensory information from the environment to make perceptual processing more efficient (Henson, 2003). In many individuals with dyslexia, however, short-term stimulus consistency appears to have a reduced effect on perception. In the auditory domain, for example, frequency discrimination thresholds in the presence of a constant reference stimulus improve for typical readers but not those with dyslexia (Ahissar et al., 2006). Individuals with dyslexia are slower to detect one of a small set of tones in noise (Chait et al., 2007) and exhibit less-accurate perception of single-talker speech in noise (Ziegler et al., 2009). Individuals with dyslexia also show impairments in learning abstract auditory categories, both natural and linguistic (e.g., voices; Perea et al., 2014; Perrachione et al., 2011) and artificial and nonlinguistic (Gabay & Holt, 2015), suggesting that deficits in short-term perceptual facilitation may be related to difficulties developing long-term perceptual representations. In the visual domain, individuals with dyslexia also have impairments in behaviors that rely upon the extraction of regularities in the sensory environment: They show elevated perceptual thresholds for sinusoidal gratings presented in noise (Sperling et al., 2005) and impaired statistical learning for both simple (Sigurdardottir et al., 2017) and complex visual stimuli (Arciuli & Simpson, 2012). Relative insensitivity to repetition and co-occurrence statistics in dyslexia may ultimately hinder the formation of abstract phonological and orthographic representations that support fluent reading (Chandrasekaran et al., 2009; Harm & Seidenberg, 2004; Seidenberg & McClelland, 1989).

An index of how regularities in the sensory environment may affect perception is the phenomenon of neural *repetition suppression*. Sometimes also called *neural adaptation*, repetition suppression describes a reduction in the neural response magnitude to repeated presentations of a stimulus (Grill-Spector et al., 2006). Neural repetition suppression has been correlated with behavioral priming, measured as faster reaction times, reduced perceptual thresholds, and better implicit memory for previously-encountered items (Schacter & Buckner, 1998). Stimulus repetition may enhance performance by attenuating the contributions of weakly-responding units to a given stimulus (Desimone, 1996) or by increasing neural synchrony (Brunet et al., 2014; Hansen & Dragoi, 2011), leading to perceptual representations that are more efficient (Wiggs & Martin, 1998) and more robust to noise (Atiani et al., 2009; Khalighinejad et al., 2019). Studies of dyslexia have observed atypical neural adaptation processes, indexed by reduced adaptation of the hemodynamic response to repetitions of voices, auditory words, and visual words, objects, and faces (Perrachione et al., 2016) and by smaller magnitude and shorter duration of neural adaptation to auditory tones (Jaffe-Dax et al., 2018; Peter et al., 2019). These differences suggest that perceptual facilitation by stimulus consistency may be impeded in dyslexia due to dysfunction in one or more of the mechanisms of rapid neural plasticity that lead to repetition suppression.

The sources of repetition suppression can be accounted for by considering perception within a neurocomputational framework for *predictive coding* (Friston, 2009; Rao & Ballard, 1999). In such a framework, sensory inputs are processed in the context of top-down predictions; unpredicted sensations (“errors”) propagate up the hierarchy in order to update those predictions (Clark, 2013). The neural response to novelty or surprise – the *prediction error* – is the learning signal that refines longer-term representations. Conversely, repetition increases predictability, thereby reducing prediction error and the concomitant neural response. Moreover, the magnitude of repetition suppression is greater when repetition is expected (Summerfield et al., 2008; Summerfield et al., 2011; Todorovic et al., 2011; Todorovic et al., 2012), implicating the involvement of top-down processes in perception that track probability, integrate over longer timescales, and establish predictions. Thus, changes in the neural response following stimulus repetition reflect the combined effect of both feedforward (bottom-up) and feedback (top-down) processing. This raises the question of whether it is differences in feedforward, feedback, or both processes that underlie the pattern of neural and behavioral repetition deficits in dyslexia.

### 1.2 The present study

While brain imaging studies have shown reduced neural adaptation to stimulus repetition in dyslexia, these prior methods have not been suitable for understanding the source of such reduction – specifically, whether it is due to differences in feedforward effects of repetition suppression, feedback effects of expectation, or both. In this study, we aimed to determine which mechanisms are responsible for reduced neurophysiological adaptation to stimulus consistency in dyslexia compared to individuals with typical reading abilities. To ascertain the relative disruption of feedforward vs. feedback signals responsible for neural adaptation in dyslexia, we measured EEG to stimulus repetition in contexts in which repetition was highly probable (and where adapted neural responses would reflect top-down expectations) or relatively improbable (and where adapted neural responses would reflect primarily feedforward repetition suppression). We considered this design in the context of three different hypotheses regarding the source of reduced perceptual and neural adaptation in dyslexia.

*Hypothesis 1: The Expectation-Deficit Hypothesis.* It may be the case that, in dyslexia, the brain fails to generate appropriate top-down expectation signals when context makes stimulus predictions possible. If so, then experimentally manipulating perceptual expectations will have little effect on the brain state of individuals with dyslexia. Expectation and attention are coupled phenomena: Cues that orient attention also activate predictions of expected stimulus features based on prior knowledge (Kastner et al., 1999; Kok, Failing, & de Lange, 2014; Summerfield & Egner, 2009). Thus, a difference in the neural correlates of expectation might indicate a relative weakness in marshaling top-down resources to organize and facilitate perception in dyslexia. This hypothesis follows from theories that emphasize a causal role for attentional deficits in dyslexia (e.g., Facoetti et al., 2000; Vidyasagar & Pammer, 2010).

*Hypothesis 2: The Feedforward-Deficit Hypothesis*. Alternatively (or additionally), it may be the case that, in dyslexia, the brain is modulated less by short-term experience. If so, then stimulus repetitions will yield less feedforward repetition suppression in dyslexia, in particular when repetition of a stimulus occurs unexpectedly, thus minimizing the top-down contributions to processing it. How could stimulus repetition not lead to a reduction in neural response, when the same population of neurons should be responsible for encoding it each time (e.g., Marlin, Hasan, & Cynader, 1998)? If repetitions of an identical stimulus are encoded in a variable manner, short-term perceptual constancy will be diminished, and its concomitant neural repetition suppression would presumably be attenuated. There is mounting evidence for the sort of neural response variability in dyslexia that could obfuscate the neural signature of repetition suppression (e.g., Centanni et al., 2018; Chandrasekaran et al., 2009; Hornickel & Kraus, 2013; Ziegler et al., 2009). This hypothesis follows from theories positing that variability and inconsistency in feedforward sensory processing are at the core of dyslexia – particularly, the *neural noise hypothesis* of dyslexia, which formalizes a model of neural hyperexcitability and stochasticity (Hancock, Pugh, & Hoeft, 2017).

*Hypothesis 3: The Expectation Integration-Deficit Hypothesis*. Finally, it may be the case that, in dyslexia, intact expectation signals are not effectively integrated into intact feedforward processing. If so, then manipulations of expectation will have similar effects on the anticipatory brain states of individuals with dyslexia as those of typical readers, but dyslexics’ neural responses to subsequent stimuli will reflect neither the reduction in prediction error that comes with fulfilled expectations (i.e., *expectation suppression;* Todorovic et al., 2011) nor the increase in prediction error triggered by a violation of expectation. This response profile has not yet been explicitly examined in dyslexia, although it may be related to reduced mismatch negativity (MMN) findings in dyslexia (Maurer et al., 2003; Neuhoff et al., 2012; Schulte-Körne et al., 2001), as properties of the MMN are well accounted for under a predictive coding framework (Baldeweg, 2007; Garrido et al., 2009; Wacongne et al., 2012). However, like the MMN-eliciting oddball paradigm, prior studies showing neural adaptation deficits in dyslexia have relied on highly predictable stimulus repetition, which precludes the ability to disentangle the effects of automatic, feedforward repetition from those of feedback-mediated stimulus expectation and prediction. This hypothesis – that intact perceptual processing and representations may be less susceptible to top-down influences in dyslexia – follows from theories suggesting that perceptual representations are intact in this disorder, but that access to them during tasks is impeded (Boets et al., 2013; Ramus & Szenkovits, 2008).

We designed the present study to adjudicate among these three hypotheses. We recorded scalp electroencephalography (EEG) from adults with and without dyslexia as they viewed stimuli under two different conditions that manipulated the expectation of stimulus repetition. We used visual stimuli because they have been shown to yield event-related potential (ERP) and spectral power effects related to manipulations of repetition and expectation (Summerfield et al., 2011). We also investigated whether the hypothesized prediction error or repetition-suppression impairments are perceptually domain-specific vs. general by using two categories of complex visual stimuli: human faces and words, each of which is processed in highly-specialized, category-specific regions of occipitotemporal cortex (Kanwisher, McDermott, & Chun, 1997; McCandliss, Cohen, & Dehaene, 2003). If adaptation effects are specific to reading or print, we would expect to see group differences for conditions involving words, but not faces (cf. Sigurdardottir et al., 2018), which could be attributed to a core dysfunction in the neural processes involved in reading. Alternatively, if adaptation effects are domain general, we would expect to see group differences in both word and face conditions, suggesting that dysfunctional neural mechanisms for prediction and adaptation are perceptually nonspecific in dyslexia (consistent with the fact that cortical processing of text is an orchestration of brain areas evolved for other purposes; Dehaene & Cohen, 2007; Price & Devlin, 2011).

Following Summerfield and colleagues (2008; 2011), we presented pairs of stimuli during experimental conditions that orthogonally manipulated the expectation of stimulus repetition vs. the stimulus repetition itself. We investigated top-down effects of ***expectation*** (Hypothesis 1) operationalized as the differential modulation of pre-stimulus spectral power between conditions with high vs. low probability of upcoming stimulus repetition. We investigated feedforward effects of ***repetition*** (Hypothesis 2) operationalized as the difference in ERPs evoked by repeated vs. novel stimuli when participants did not expect stimuli to repeat. Finally, we investigated the integrated top-down/feedforward effects of ***prediction error*** (Hypothesis 3) operationalized as the difference between ERPs elicited by stimuli that fulfilled vs. violated the expectation of repetition. By determining how individuals with dyslexia differ from typical readers in these three components of predictive processing, we can ascertain whether behavioral and neural adaptation deficits in dyslexia are attributable to differences in bottom-up mechanisms of repetition suppression, top-down mechanisms of expectation and prediction error, or their integration.

## 2. Methods

### 2.1 Participants

Individuals with dyslexia (*N* = 20; 12 female, 8 male; age 19–32 years, mean ± standard deviation = 25 ± 4) and typical readers (*N* = 20; 9 female, 11 male; age 18–31 years, 23 ± 4) participated in this study. All were native speakers of English who reported no history of neurological disorder. All participants scored 90 or above on the Performance IQ subscale of the Wechsler Abbreviated Scale of Intelligence (WASI; Wechsler, 1999). Inclusionary criteria for the Dyslexia group consisted of a prior clinical diagnosis or lifelong history of reading impairment, in addition to current, age-based standard scores of 90 or below on two or more of the following four measures: Word Identification and Word

Attack subtests from the Woodcock Reading Mastery Tests (WRMT; Woodcock, 1998) and Sight Word Efficiency and Phonemic Decoding Efficiency subtests from the Test of Word Reading Efficiency (TOWRE; Torgesen, Wagner, & Rashotte, 1999). Members of the typical-reader (Control) group scored above 90 on each of those four measures and reported no history of reading or language difficulties.

Psychometric characterization of the two groups is summarized in **Table 1**. Informed, written consent was obtained from all participants, as approved and overseen by the Massachusetts Institute of Technology Committee on the Use of Humans as Experimental Subjects.

**Table 1.** Behavioral characterization of the Control and Dyslexia groups.

| Test | Subtest | Control |  | Dyslexia |  | Difference |  |  |
| --- | --- | --- | --- | --- | --- | --- | --- | --- |
|  |  | Mean | SD | Mean | SD | *t* | *p* | *d* |
| WASI | Performance IQ | 118.95 | 8.82 | 110.90 | 10.94 | 2.56 | 0.01 | 0.81 |
| WAIS-IV | Digit Span Total | 11.75 | 2.92 | 9.35 | 2.50 | 2.79 | 0.008 | 0.88 |
| CTOPP | Elision | 11.20 | 0.83 | 8.50 | 2.48 | 4.61 | 0.0001 | 1.46 |
|  | Blending Words | 12.35 | 1.87 | 11.00 | 2.20 | 2.09 | 0.04 | 0.66 |
|  | Nonword Repetition | 9.90 | 3.06 | 8.15 | 1.18 | 2.39 | 0.03 | 0.75 |
| WRMT-R/NU | Word Identification | 109.35 | 8.32 | 94.05 | 7.13 | 6.24 | < 0.0001 | 1.97 |
|  | Word Attack | 108.40 | 11.80 | 92.00 | 6.85 | 5.37 | < 0.0001 | 1.70 |
| TOWRE | Sight Word Efficiency | 107.70 | 8.47 | 84.65 | 7.69 | 9.01 | < 0.0001 | 2.85 |
|  | Phonemic Decoding Efficiency | 108.25 | 7.43 | 79.30 | 6.59 | 13.04 | < 0.0001 | 4.12 |
| WJIII ToA | Reading Fluency | 126.20 | 13.58 | 98.00 | 13.35 | 6.54 | < 0.0001 | 2.09 |

**Abbreviations:** WASI: *Wechsler Abbreviated Scale of Intelligence* (Wechsler, 1999); WAIS-IV: *Wechsler Adult Intelligence Scale – Fourth Edition* (Wechsler, 2008); CTOPP: *Comprehensive Test of Phonological Processing* (Wagner, Torgesen, & Rashotte, 1999); WRMT-R/NU: *Woodcock Reading Mastery Tests – Revised/Normative Update* (Woodcock, 1998); TOWRE: *Test of Word Reading Efficiency* (Torgesen, Wagner, & Rashotte, 1999); WJIII ToA: *Woodcock-Johnson III Test of Achievement* (Woodcock, McGrew, & Mather, 2001). The Reading Fluency score is missing for one participant in the Dyslexia group.

### 2.2. Stimuli

Face stimuli consisted of 660 unique color photographs of front-facing men and women with neutral expressions positioned against black backgrounds, taken from collections such as the Karolinska Directed Emotional Faces, the NimStim Face Stimulus Set, and the Radboud Faces Database^1^. Word stimuli were 660 unique monosyllabic English nouns (e.g., *boon*, *sled*, *wheat*) written in lowercase Arial typeface and presented in black on a white background.

### 2.3 Procedure

**Figure 1** illustrates the experimental task design. Participants performed a rare target-detection task while viewing pairs of visually presented faces or words. Across trials, we varied whether the second stimulus (***S2***) was a repeat of the first (***S1***); across blocks, we varied the probability of such a repetition. Face and Word stimuli were presented in two separate runs, each lasting ~30 minutes. Five, three-minute blocks each of the Expect Repeat and Expect Change conditions alternated throughout each run, with visual instructions preceding each: “Now you will see the repeating condition. You will usually see each face twice in a row. Watch for the upside-down faces,” or, “Now you will see the non-repeating condition. You will usually see each face only once. Watch for the upside-down faces,” (or “words”), respectively. In the Expect Repeat condition, Expected Repetitions occurred on 75% of trials and Unexpected Changes occurred on 25% of trials; in the Expect Change condition, these probabilities were reversed, with 25% Unexpected Repetition and 75% Expected Change trials (Summerfield et al., 2008; Kaliukhovich & Vogels, 2010). Prior to the experiment, participants were given an oral explanation of the two conditions in the same language quoted above.

**Figure 1.**
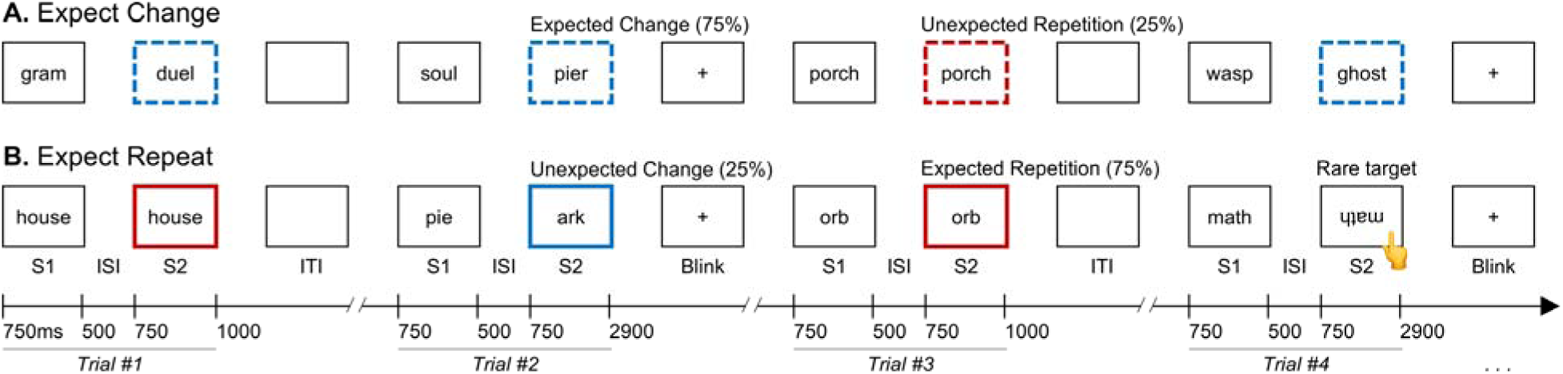
Task design. Participants viewed pairs of stimuli under conditions that manipulated the probability of stimulus repetition. Each trial consisted of a pair of stimuli, S1 and S2. **(A)** In the Expect Change condition, participants were t ld to expect to see each stimulus only once. The S2 stimulus differed from the S1 stimulus on 75% of trials (*Expected Changes*) and was the same as the S1 stimulus on 25% of trials (*Unexpected Repetitions*). **(B)** In the Expect Repeat condition, participants were told to expect to see each stimulus twice in a row. The S2 stimulus was the same as the S1 stimulus on 75% of trials (*Expected Repetitions*) and differed from the S1 stimulus on 25% of trials (*Unexpected Changes*). In all conditions, participants pressed a button whenever they observed an upside-down stimulus (illustrated in Trial #4 above), which occurred on approximately 5% of trials. For brevity, only the Words condition is shown; the trial structure was identical in the Faces condition. The convention of line colors and dashing denoting conditions is consistent with Figures 4 and 5.

Each stimulus (S1 and S2) was presented for 750 ms, with a 500-ms inter-stimulus interval (ISI) between stimuli in a pair. The screen was blank for 500 ms before and after each trial (an effective 1000 ms inter-trial interval (ITI). After every two trials, a fixation cross appeared on the screen for 1900 ms, which served as participants’ cue to blink. Participants were asked to refrain from blinking until the blink cue. Approximately one minute of practice was administered in order to familiarize participants with the procedure and with the timing of the blink cue. To ensure attention, participants performed the target-detection task by pressing a button with their right hand in response to any upside-down face or word. These targets appeared on ~5% of trials, distributed pseudorandomly across conditions. Targets always appeared on the S2 stimulus. Trials containing targets were analyzed for participants’ behavioral responses (i.e., response time and accuracy) but discarded from electrophysiological analyses. Each participant was exposed to 440 trial pairs during the Faces run and 440 trial pairs during the Words run, with the run order balanced within and across groups. To avoid item-specific effects, each participant viewed one of four counterbalanced stimulus lists for Faces and for Words. No word or face stimulus appeared in more than one trial per participant.

### 2.4 EEG data acquisition

The experiment was conducted in a sound-attenuating, electrically-shielded booth in which participants were seated in front of a cathode-ray tube monitor with a 60-Hz refresh rate. EEG was recorded during the task with the Biosemi ActiveTwo System (Biosemi, Amsterdam), using a 32-electrode cap conforming to the international 10-20 system. External electrodes were placed on the left and right mastoids and the tip of the nose, and electro-oculograms were recorded from the left infra-orbital ridge and the right lateral canthus. Impedance was ensured to be <40 µV in each channel. EEG was recorded with a low-pass hardware filter with a half-power cutoff at 104 Hz and digitized at 512 Hz with 24 bits of resolution.

### 2.5 Behavioral data analysis

In order to characterize differences between the Control and Dyslexia groups in reading, phonological, or cognitive measures, we compared standard scores on the behavioral assessments using independent-samples *t*-tests, and computed Cohen’s *d* as a measure of effect size. (Note that performance on the WRMT and TOWRE were the inclusionary criteria, and group differences should be interpreted accordingly.)

To determine whether the groups differed in processing the stimuli during EEG recording, we compared their response times and accuracy (percent of targets detected) in two repeated-measures analyses of variance (ANOVA) with within-subjects factors of *Stimulus* (Faces or Words) and *Condition* (Expect Repeat or Expect Change), and a between-subjects factor of *Group* (Control or Dyslexia). All tests were performed in R 3.3.3 (R Core Team, 2017).

### 2.6 EEG/ERP data analysis

#### 2.6.1 Preprocessing

A separate EEG dataset was created for each stimulus type (i.e., Faces and Words). The data were preprocessed using the EEGLAB 12.0.2.6b toolbox (Delorme & Makeig, 2004) in MATLAB 2014a (The MathWorks, Natick, MA). Data were referenced to the average of the left and right mastoids and band-pass filtered from 0.1 to 100 Hz using a zero-phase, windowed-sinc, finite-impulse-response filter. Continuous data were epoched from –1000 to 3150 ms with respect to the onset of each stimulus, excluding the rare upside-down target stimuli. These epochs captured the interval during which participants were instructed to blink. Independent components analysis was performed and components whose spectra and scalp topography were characteristic of blinks, muscle artifact, or single-trial electrode pops were removed from the data. On average, across participants, 3.69 (s.d. = 1.83, range = 1–10) of 32 components were removed from each dataset. Subsequently, any epoch in which the peak-to-peak voltage exceeded 200 μV was removed from the dataset. Of the two frequent trial types (Expected Repetitions and Expected Changes), an average of 151.18 trials (s.d. = 7.49, range = 107–164) remained after artifact rejection; this number did not differ by group or stimulus type (repeated-measures ANOVA; all *F’*s < 0.42, *p’s* > 0.52). Of the two infrequent trial types (Unexpected Repetitions and Unexpected Changes), an average of 53.31 trials (s.d. = 2.63, range = 36–57) remained; this number did not differ by group or stimulus type (repeated-measures ANOVA; all *F’*s < 0.52, *p’s* > 0.47).

#### 2.6.2 Time-frequency representations

Time-frequency representations (TFRs) of single-trial EEG were calculated with a multi-taper convolution method, with a Hanning taper for frequencies from 1 to 20 Hz in 1-Hz steps using the *ft_freqanalysis* function in the FieldTrip toolbox (version 18-02-22; Oostenveld et al., 2011). The window length was fixed at 1000 ms, sliding by 50-ms steps from –500 to 2500 ms relative to the onset of the first stimulus (S1); times reported in the Results section indicate the center timepoint of a 50-ms analysis window. The single-trial TFR data were baseline corrected by calculating the relative power change with respect to the average power estimate from –500 to 0 ms (prior to S1 onset) over all trials (all conditions).

As the TFRs index the neural oscillatory power reflecting differences in ongoing brain state, we focused on the TFRs prior to the presentation of the second stimulus (S2) in conditions that modulated participants’ ***expectation*** that S2 would repeat (Expect Repeat) or change (Expect Change) from S1. To understand the effects of expectation on neural oscillatory power in anticipation of S2, we used data from the Control group to identify data-driven regions of interest (ROIs) during the time window from 0 to 1250 ms relative to S1 onset. In the Faces and Words conditions separately, we contrasted the TFRs from all trials in the Expect Repeat condition with those from all trials in the Expect Change condition, regardless of whether the trial went on to be a Repetition or a Change at S2. Specifically, at the subject level, we computed the differences in mean oscillatory power in the Expect Repeat vs. the Expect Change condition for each time × channel × frequency data point. At the group level, the subject-level mean differences across frequencies (1–20 Hz), all 32 scalp channels, and time (0–1250 ms relative to S1 onset) were contrasted against zero with a dependent-samples *t*-test (Maris & Oostenveld, 2007). We employed cluster-based correction based on the Monte Carlo significance probability estimated with 2000 random partitions. This procedure resulted in frequency × channel × time clusters that revealed significant differences in the Expect Repeat and Expect Change conditions (two-tailed *α* < 0.05). For each expectation cluster and for each participant, we extracted the mean TFR value within the cluster, averaged across trials, for the Expect Repeat and Expect Change conditions. We submitted these values to a repeated-measures ANOVA that included a within-subject factor of *Expectation* (Expect Repeat vs. Expect Change) and a between-subjects factor of *Group* (Control vs. Dyslexia). Post-hoc *t*-tests were performed on significant interactions. This analysis tested whether effects of perceptual expectation differed by group (Hypothesis 1).

#### 2.6.3 Event-related potentials

Event-related potentials (ERPs) were calculated using the *ft_timelockanalysis* function in FieldTrip. The single-trial data were baseline corrected by subtracting the mean voltage over the 200 ms immediately preceding trial onset.

As the ERPs index the rapid neural response to a stimulus ***repetition*** or change, we focused on the ERPs during the presentation of the second stimulus (S2) that changed from vs. repeated the first stimulus (S1). In order to examine whether and how these neural responses differed between the Dyslexia and Control groups, we first defined data-driven ROIs based on the evoked responses of the Control-group participants. In the Faces and Words conditions separately, we contrasted evoked responses to S2 that changed from vs. repeated S1. We defined two different types of ERP ROIs, *repetition* ROIs and *prediction error* ROIs:

***Repetition ROIs*** were defined in the Expect Change condition, when stimulus-specific predictability was low. The difference between the frequent Expected Change trials and the infrequent Unexpected Repetition trials represents the neural signature of feedforward repetition effects (Expected Change – Unexpected Repetition).

***Prediction-error ROIs*** were defined in the Expect Repeat condition, when stimulus-specific predictability was high. The frequent Expected Repetition trials represent fulfilled expectations, whereas the infrequent Unexpected Change trials represent violated expectations; therefore, the difference in neural response magnitude between these trial types indexes the neural signature of prediction error (Unexpected Change – Expected Repetition).

Specifically, for the Expect Repeat and Expect Change conditions separately, we computed single-subject-level ERP differences between Change and Repetition trials during the presentation of S2. Across subjects in the Control group, the mean ERP differences across all 32 scalp channels and the 750 ms between the onset and offset of S2 were compared against zero via a permutation-based dependent samples *t*-test. As in the TFR analysis, the significance of a cluster was based on 2000 random Monte Carlo iterations. Channel × time clusters identified by this procedure indicate that Change and Repetition trials are significantly different (two-tailed *α* < 0.05).

Within each ***repetition*** cluster, we extracted the mean ERP to S2 in the Unexpected Repetition and Expected Change trials. We submitted these values from each participant to a repeated-measures ANOVA with a within-subject factor of *Repetition* (Repetition vs. Change) and a between-subjects factor of *Group* (Control vs. Dyslexia). Post-hoc *t*-tests were performed on significant interactions. This analysis tested whether feedforward repetition effects differed by group (Hypothesis 2).

Within each ***prediction-error*** cluster, we extracted the mean ERP to S2 for each of the four trial types (Expected Repetitions, Unexpected Changes, Unexpected Repetitions, and Expected Changes). We submitted these values from each participant to a repeated-measures ANOVA that included within-subject factors for the experimental manipulations (*Repetition*: Repetition vs. Change; *Expectation*: Expect Repeat vs. Expect Change) and a between-subjects factor of *Group* (Control vs. Dyslexia). This analysis examined whether expectation affected the evoked neural response to repetition differently across groups (Hypothesis 3). Note that our approach of first defining ROIs by Change vs. Repetition and then testing for any orthogonal interaction with expectation avoids circular inference (Summerfield et al., 2011). Post-hoc paired- or independent-sample *t*-tests (as appropriate) were performed to explore significant interactions, and Cohen’s *d* was calculated as a measure of effect size. Statistical analyses were performed with the *ez* and *effsize* packages in R.

Finally, to allow for the possibility that the Control and Dyslexia groups might show equivalent or even divergent effects of expectation, repetition, and/or prediction error at different topographies, latencies, or frequencies, we repeated the analyses described above, this time first defining each type of ROI based on data from the Dyslexia group, and then analyzing between-group differences as described above.

## 3. Results

### 3.1 Behavioral results

By design, the Dyslexia group performed significantly below the Control group on the four measures of single-word reading serving as inclusionary criteria; they also performed significantly below the Control group on measures of phonological processing, sentence-reading fluency, digit span, and performance IQ (**Table 1**).

Participants performed with high accuracy on the incidental target detection task during EEG recording, as detailed in **Table 2**. There were no significant differences in accuracy or response time between the Control and Dyslexia groups, and no significant effects of stimulus type or expectation condition (accuracy: all *F*’s < 2.02, *p*’s > 0.16; response time: all *F*’s < 2.25, *p*’s > 0.14).

**Table 2.**
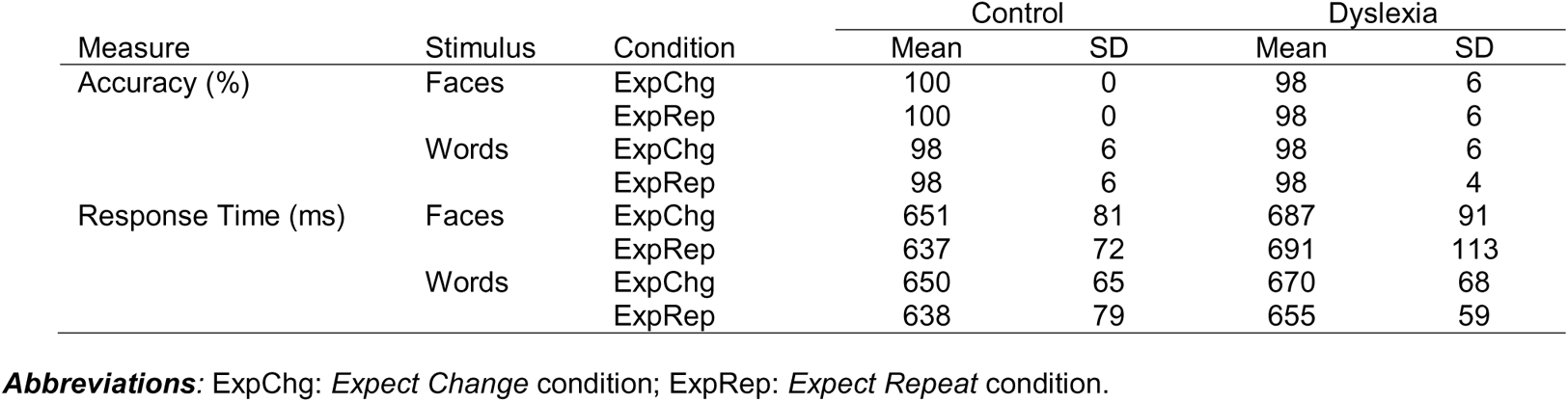
Target detection task performance.

| Measure | Stimulus | Condition | Control |  | Dyslexia |  |
| --- | --- | --- | --- | --- | --- | --- |
|  |  |  | Mean | SD | Mean | SD |
| Accuracy (%) | Faces | ExpChg | 100 | 0 | 98 | 6 |
|  |  | ExpRep | 100 | 0 | 98 | 6 |
|  | Words | ExpChg | 98 | 6 | 98 | 6 |
|  |  | ExpRep | 98 | 6 | 98 | 4 |
| Response Time (ms) | Faces | ExpChg | 651 | 81 | 687 | 91 |
|  |  | ExpRep | 637 | 72 | 691 | 113 |
|  | Words | ExpChg | 650 | 65 | 670 | 68 |
|  |  | ExpRep | 638 | 79 | 655 | 59 |

**Abbreviations:** ExpChg: *Expect Change* condition; ExpRep: *Expect Repeat* condition.

### 3.2 Effects of perceptual expectations on neural oscillations

In order to evaluate Hypothesis 1 (group differences in generating top-down perceptual expectations), we investigated whether manipulating participants’ expectations about stimulus repetition between the Expect Repeat and Expect Change conditions translated into modulations of spectral power prior to S2 presentation, and whether this modulation differed between Control and Dyslexia groups.

#### 3.4.1 Faces

As defined in the Control group, the EEG was significantly modulated by expectation. One broadly-distributed cluster, observed spectrally from 6 to 13 Hz and temporally from 100 to 1250 ms (with respect to S1 onset), was identified as sensitive to *Expectation*, showing oscillatory desynchronization in the Expect Repeat condition relative to the Expect Change condition (**Figure 2A-C**). Within this cluster there was no main effect of *Group*, nor a *Group* × *Expectation* interaction (**Table 3**).

**Figure 2.**
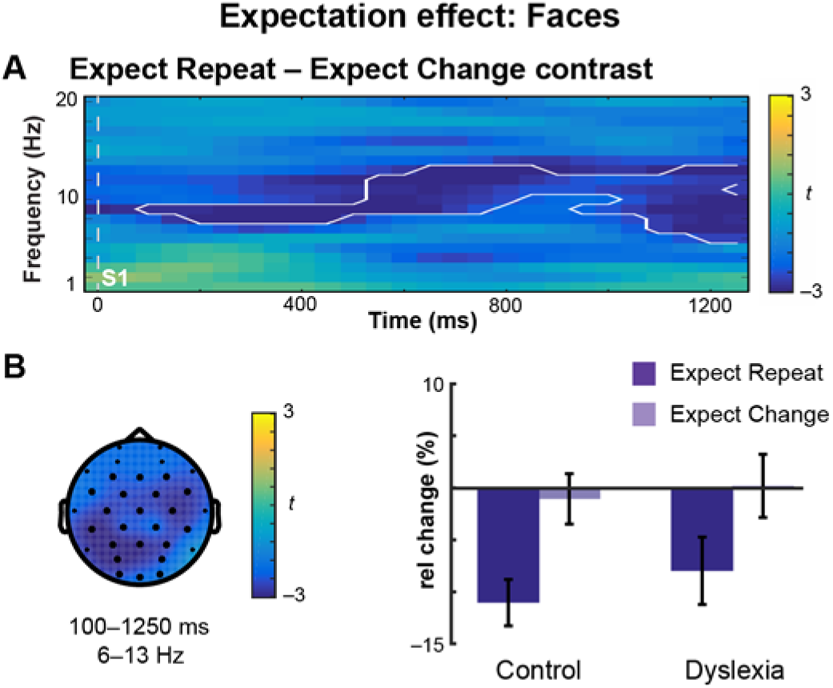
Expectation of face repetition versus change modulates neural oscillations. **(A)** Oscillatory power prior to S2 onset plotted in Control subjects as a time-frequency representation of the Expect Repeat – Expect Change contrast, where color indicates the *t*-statistic. One extended cluster (white outline) was identified in the Control group. **(B)** Topographical plot of the cluster. The *t*-statistic is averaged over time and frequency. Dark electrodes belong to the cluster. Barplot of the mean-difference values extracted from the cluster for Control and Dyslexia groups, expressed as a percent change from baseline. Error bars represent (between-subjects) SEM. Both groups showed desynchronization in the Expect Repeat condition.

**Table 3.** Expectation effects.

| Cluster | Time (ms) | Freq. (Hz) | Distribution | Polarity | ANOVA Results | $F_{1,38}$ | $p$ | |
| --- | --- | --- | --- | --- | --- | --- | --- | --- |
| **Faces** |  |  |  |  |  |  |  |  |
| #1 | 100–1250 | 6–13 | global | ExpChg > ExpRep | Group | 0.34 | 0.6 |  |
|  |  |  |  |  | Expectation | 51.61 | < 0.0001 | *** |
| | | | | | Group × Expectation | 0.52 | 0.5 | |
| **Words** |  |  |  |  |  |  |  |  |
| #1 | 0–150 | 10 | central | ExpChg > ExpRep | Group | 1.01 | 0.3 |  |
|  |  |  |  |  | Expectation | 18.69 | 0.0001 | *** |
| | | | | | Group × Expectation | 0.01 | 0.9 | |
| #2 | 0–1150 | 6–8 | global | ExpChg > ExpRep | Group | 1.13 | 0.3 |  |
|  |  |  |  |  | Expectation | 43.85 | < 0.0001 | *** |
| | | | | | Group × Expectation | 5.11 | 0.03 | * |

Cluster time ranges are given with respect to the onset of S1. **Abbreviations** as in Table 2.

As defined in the Dyslexia group, the EEG also distinguished expectation conditions. Three clusters covering several alpha frequencies (8–9 and 13 Hz) showed desynchronization in the Expect Repeat condition relative to the Expect Change condition; there were again no main effects of nor interactions with *Group* (**Supp. Table 1**).

#### 3.4.2 Words

In the Control group, two clusters were identified as sensitive to expectation, one in the alpha range (Cluster #1; 10 Hz; 0–150 ms), and the other in the theta to low-alpha range (Cluster #2; 6–8 Hz; 0–1150 ms) (**Figure 3A**; **Table 3**). In the alpha cluster (#1), the effect of *Expectation* was characterized by desynchronization in the Expect Repeat condition in both groups (**Figure 3B**). Within the theta cluster (#2) there was a significant *Group* × *Expectation* interaction; post-hoc tests indicated that theta oscillations in both the Control group (*t* = 6.57; *p* < 0.0001; *d* = 1.09) and the Dyslexia group (*t* = 2.96; *p* = 0.008; *d* = 0.47) were modulated by expectation condition, showing synchronization in the Expect Change condition relative to the Expect Repeat condition (**Figure 3C**). Theta power did not significantly differ between the groups in the Expect Change condition (*t* = −0.16; *p* = 0.9; *d* = 0.05) or the Expect Repeat condition (*t* = −1.87; *p* = 0.07; *d* = 0.59).

**Figure 3.**
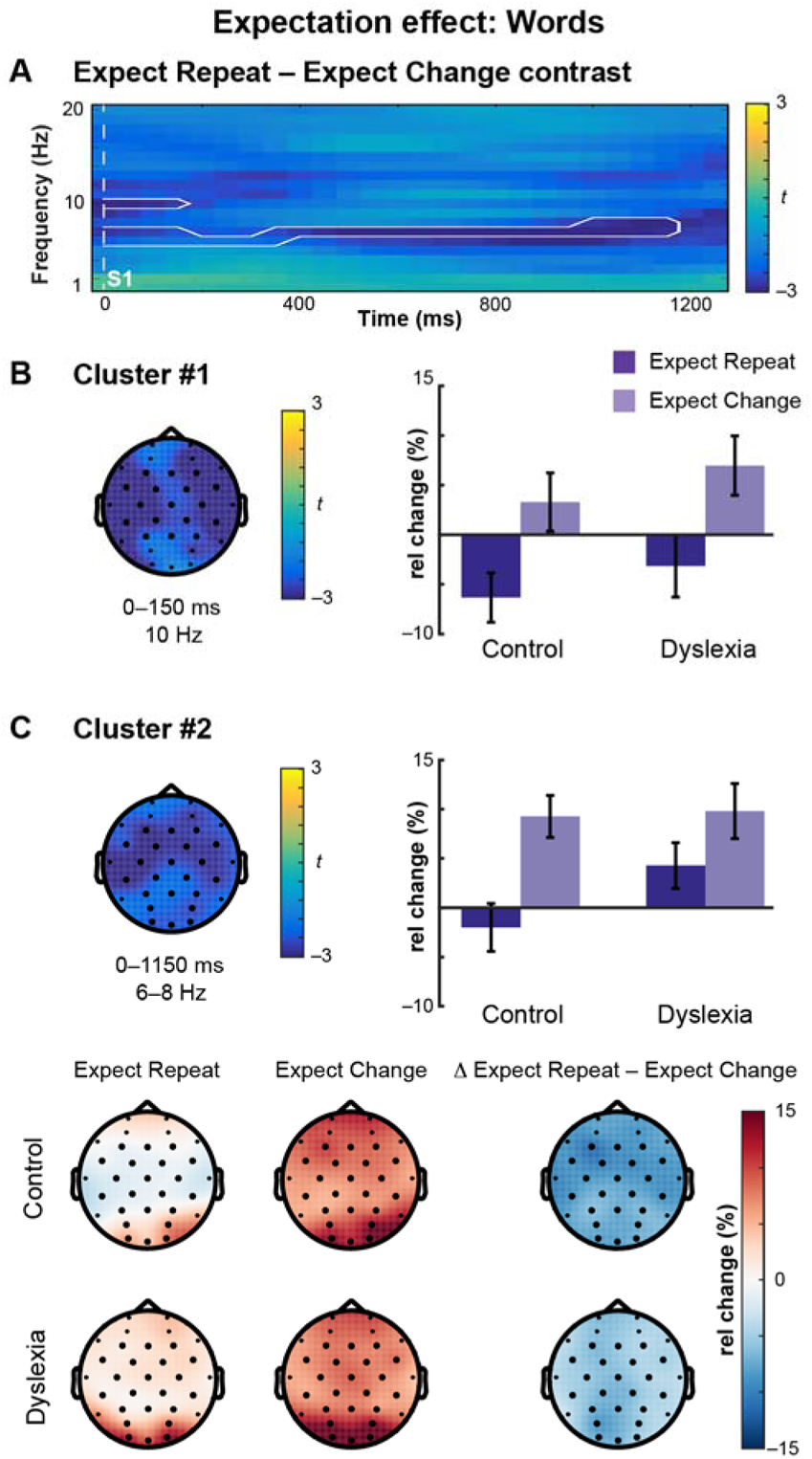
Expectation of word repetition versus change modulates neural oscillations. **(A)** Oscillatory power prior to S2 onset plotted in Control subjects as a time-frequency representation of the Expect Repeat – Expect Change contrast, where color indicates the *t*-statistic. Two clusters (white outlines) were identified in the Control group. **(B)** Topographical plot of the cluster at 10 Hz. The *t*-statistic is averaged over time and frequency. Dark electrodes belong to the cluster. In the barplot, mean-difference values extracted from the cluster for Control and Dyslexia groups demonstrate desynchronization in the Expect Repeat condition. Error bars represent (between-subjects) SEM. **(C)** Topographical plot of the cluster at 6–8 Hz. The *t*-statistic is averaged over time and frequency. Dark electrodes belong to the cluster. In the barplot, mean-difference values extracted from the cluster for Control and Dyslexia groups demonstrate synchronization in the Expect Change condition. A significant *Group* × *Expectation* interaction is plotted in detail in the lower panel. Each group showed synchronization in the Expect Change condition. Post-hoc tests revealed significant expectation condition-related modulation in each group.

In the Dyslexia group, two alpha-range expectation clusters were identified: 11–12 and 8 Hz (**Supp. Table 1**), with desynchronization observed in the Expect Repeat condition relative to the Expect Change condition. In the 8-Hz cluster, there was a *Group* × *Expectation* interaction; post-hoc tests revealed that this was driven by an *Expectation* effect present in the Dyslexia group (*t* = 4.54; *p* = 0.0002; *d* = 0.66) but not in the Control group (*t* = 0.92; *p* = 0.4; *d* = 0.15).

### 3.3 Effects of repetition on ERPs

In order to evaluate Hypothesis 2 (group differences in feedforward effects of stimulus repetition), we investigated ERPs within ROIs defined by neural responses reflecting *repetition effects* (Unexpected Repetitions vs. Expected Changes in the Expect Change condition). Hypothesis 2 is tested in the 2-way interaction (*Group* × *Repetition*), testing whether the feedforward neural signature of repetition differed between groups.

#### 3.3.1 Repetition Effects: Faces

ERPs evoked by Unexpected Repetitions vs. Expected Changes of faces were not significantly different in either the Control group (**Table 4**) or the Dyslexia group (**Supp. Table 2**).

**Table 4.** Repetition effects.

| Cluster | Time (ms) | Distribution | Polarity | ANOVA Results | $F_{1,38}$ | $p$ | |
| --- | --- | --- | --- | --- | --- | --- | --- |
| **Faces** |  |  |  |  |  |  |  |
| n/a | n/a | n/a | n/a | n/a |  |  |  |
| **Words** |  |  |  |  |  |  |  |
| #1 | 365–512 | global | Rep > Chg | *Group* | 0.07 | 0.8 |  |
|  |  |  |  | *Repetition* | 99.61 | < 0.0001 | *** |
| | | | | *Group* × *Repetition* | 0.28 | 0.6 | |
| #2 | 518–522 | central | Rep > Chg | *Group* | 0.10 | 0.8 |  |
|  |  |  |  | *Repetition* | 48.10 | < 0.0001 | *** |
| | | | | *Group* × *Repetition* | 0.11 | 0.7 | |

Clusters are numbered chronologically. Cluster time ranges are given with respect to the onset of S2 in the Expect Change condition. **Abbreviations:** Chg: *Change* trials; Rep: *Repetition* trials.

#### 3.3.2 Repetition Effects: Words

In the Control group, there was a significant difference between Expected Changes and Unexpected Repetitions. Statistical tests identified clusters at 365–512 and 518–522 ms within which there was a significant effect of *Repetition* (**Table 4**). In both clusters, the main effect of *Group* and the *Group* × *Repetition* interaction were not significant.

In the Dyslexia group, Expected Change and Unexpected Repetition trials were also significantly different, and four clusters were identified: 365–367, 379–520, 527–539, and 553–561 ms (**Supp. Table 2**). In no cluster was there a main effect of *Group*. In the third cluster a significant *Group* × *Repetition* interaction was driven by a significant *Repetition* effect in the Dyslexia group (*t* = −4.81; *p* = 0.0001; *d* = 0.53) but not in the Control group (*t* = −0.62; *p* = 0.5; *d* = 0.13).

### 3.4 Effects of prediction error on ERPs

In order to evaluate Hypothesis 3 (group differences in integration of top-down expectation and feedforward perception), we investigated ERPs within ROIs defined by neural responses reflecting *prediction error* (Unexpected Changes vs. Expected Repetitions). Hypothesis 3 is tested in a 3-way interaction (*Group* × *Repetition* × *Expectation*), testing whether the distinct neural signature of prediction error differed between groups.

#### 3.4.1 Prediction Error: Faces

Grand-average ERPs evoked by Repetitions and Changes in each expectation condition are shown for each group in **Figure 4A**. The data-driven ROI definition revealed three clusters that demonstrated significant prediction-error effects in the Control group: Cluster #1 encompasses a broad central effect at 258–305 ms, Clusters #2 and #3 encompass two posterior effects at 391–406 and 428– 440 ms, respectively (**Table 5**; **Figures 4B** (shaded regions) and **4C**).

**Figure 4.**
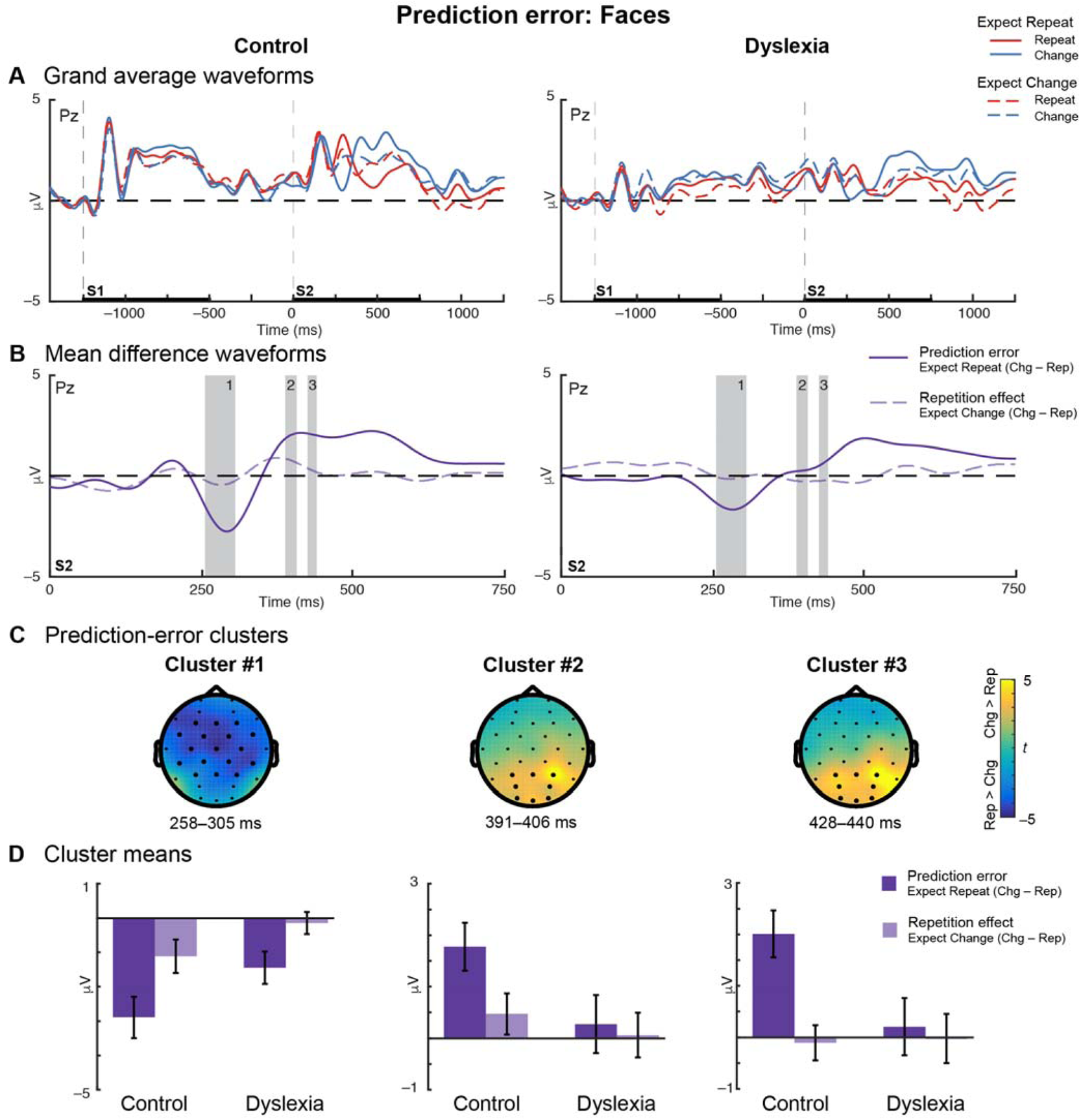
Reduced prediction error in dyslexia for unexpected changes versus expected repetitions of faces. **(A)** Grand-average waveforms for Control (left) and Dyslexia (right) groups plotted at representative electrode Pz show that ERPs diverge during the during the second stimulus (S2) interval for Repetition (red) and Change (blue) trials under the expectation of repetition (solid lines) or of change (dashed lines). **(B)** Mean-difference waveforms for prediction error in the Expect Repeat condition (solid dark purple) and repetition in the Expect Change condition (dashed light purple) during S2 presentation. Control data are plotted on the left and Dyslexia on the right; gray bars on both indicate the durations of the prediction-error clusters identified in the Control group. **(C)** Topographical plots for each of the three prediction-error clusters identified in the Control group. Color indicates the prediction-error effect expressed as a *t*-statistic, averaged over the duration of the cluster. Dark electrodes significantly differentiate Change versus Repeat trials. **(D)** Mean-difference voltage values extracted from each cluster for Control and Dyslexia groups. Error bars represent (between-subjects) SEM. Overall, greater voltage differences are observed under the Expect Repeat condition than the Expect Change condition. In each cluster, prediction error is significantly or trends larger in Control versus Dyslexia.

**Table 5.**
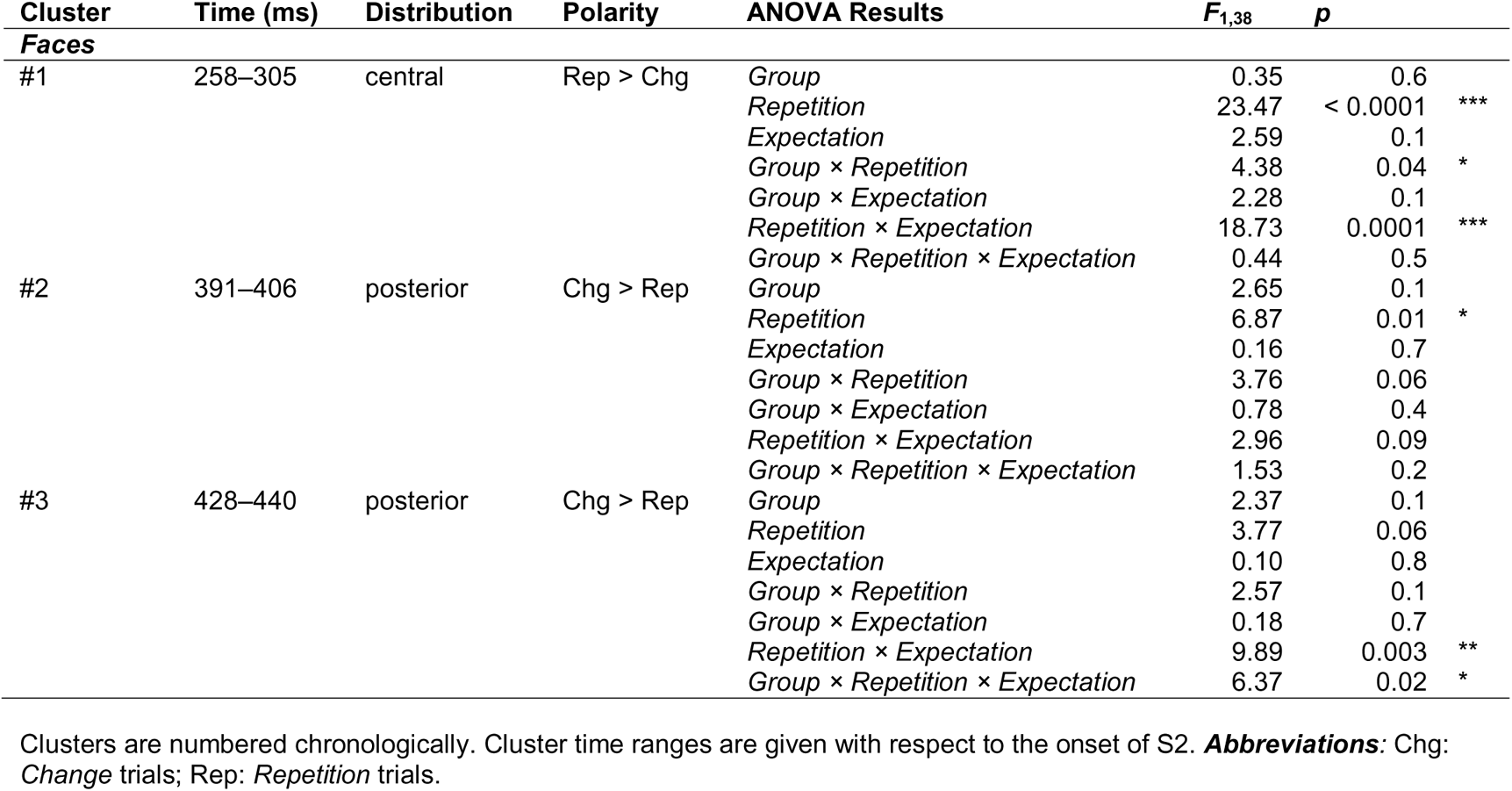
Prediction-error effects (Faces)

##### 3.4.1.1 Faces Cluster #1

Repeated-measures ANOVA performed on the mean voltages extracted from the first prediction-error cluster identified a *Group* × *Repetition* interaction (**Table 5**). Post-hoc tests revealed that, across expectation conditions, Change and Repetition trials were significantly different in both the Control group (*t* = −4.34; *p* = 0.0004; *d* = 0.75) and the Dyslexia group (*t* = −2.29; *p* = 0.03; *d* = 0.29), but the *Repetition* effect size was larger in the Control group. The separation of Change and Repetition ERPs between 258 and 305 ms can be seen in the blue vs. red traces in **Figure 4A**. Additionally, a significant *Repetition × Expectation* interaction was identified in this cluster; the magnitude of the prediction error effect was larger than that of the repetition effect (*t* = −4.36; *p* < 0.0001; *d* = 0.69), indicating that the difference between Changes and Repetitions was greater when a repetition was expected vs. unexpected (**Figures 4A** (solid vs. dashed traces) and **4D, left**).

##### 3.4.1.2 Faces Cluster #2

The second cluster showed a main effect of *Repetition* and trends toward *Repetition* × *Group* and *Repetition* × *Expectation* interactions (**Table 5**; **Figure 4D, middle**).

##### 3.4.1.3 Faces Cluster #3

The third cluster was characterized by a *Group* × *Repetition* × *Expectation* interaction (**Table 5**). Four post-hoc tests were conducted to explore this result. The prediction-error effect was significantly greater in the Control group than in the Dyslexia group *(t* = 2.51; *p* = 0.02; *d* = 0.80), while the repetition effect did not differ significantly between groups *(t* = −0.13; *p* = 0.9; *d* = 0.04). In the Control group, the prediction-error effect was significantly greater than the repetition effect (*t* = 4.09; *p* = 0.0006; *d* = 1.16); however, in the Dyslexia group, the magnitudes of these effects were not significantly different (*t* = 0.43; *p* = 0.7; *d* = 0.10). Altogether, the three-way interaction was driven by a disproportionately large prediction-error effect (relative to the feedforward repetition effect) in the Control group (**Figure 4D, right**). The mean-difference waveforms (**Figure 4B**) reveal weak repetition effects for faces throughout the epoch in both groups. Prediction-error effects are evident in an earlier window (~300 ms) at central sites, and in the opposite polarity in a later window (~400 ms) at posterior sites – for which the onset is earlier in the Control group.

##### 3.4.1.4 Clusters defined in the Dyslexia group

When the ROI definition was performed in the Dyslexia group instead of in the Control group, no significant difference between Unexpected Change and Expected Repetition ERPs was observed, and thus no prediction-error clusters for Faces were identified (**Supp. Table 3**).

#### 3.4.2 Prediction Error: Words

Grand-average ERPs evoked by word repetitions and changes in each expectation condition are shown for each group in **Figure 5A**. In the Control group, ERPs evoked by Unexpected Changes and Expected Repetitions were also significantly different. Statistical tests identified three centrally-distributed prediction-error clusters: 295–303, 307–311, and 383–395 ms (**Table 6**; **Figures 5B** (gray bars) and **5C**).

**Figure 5.**
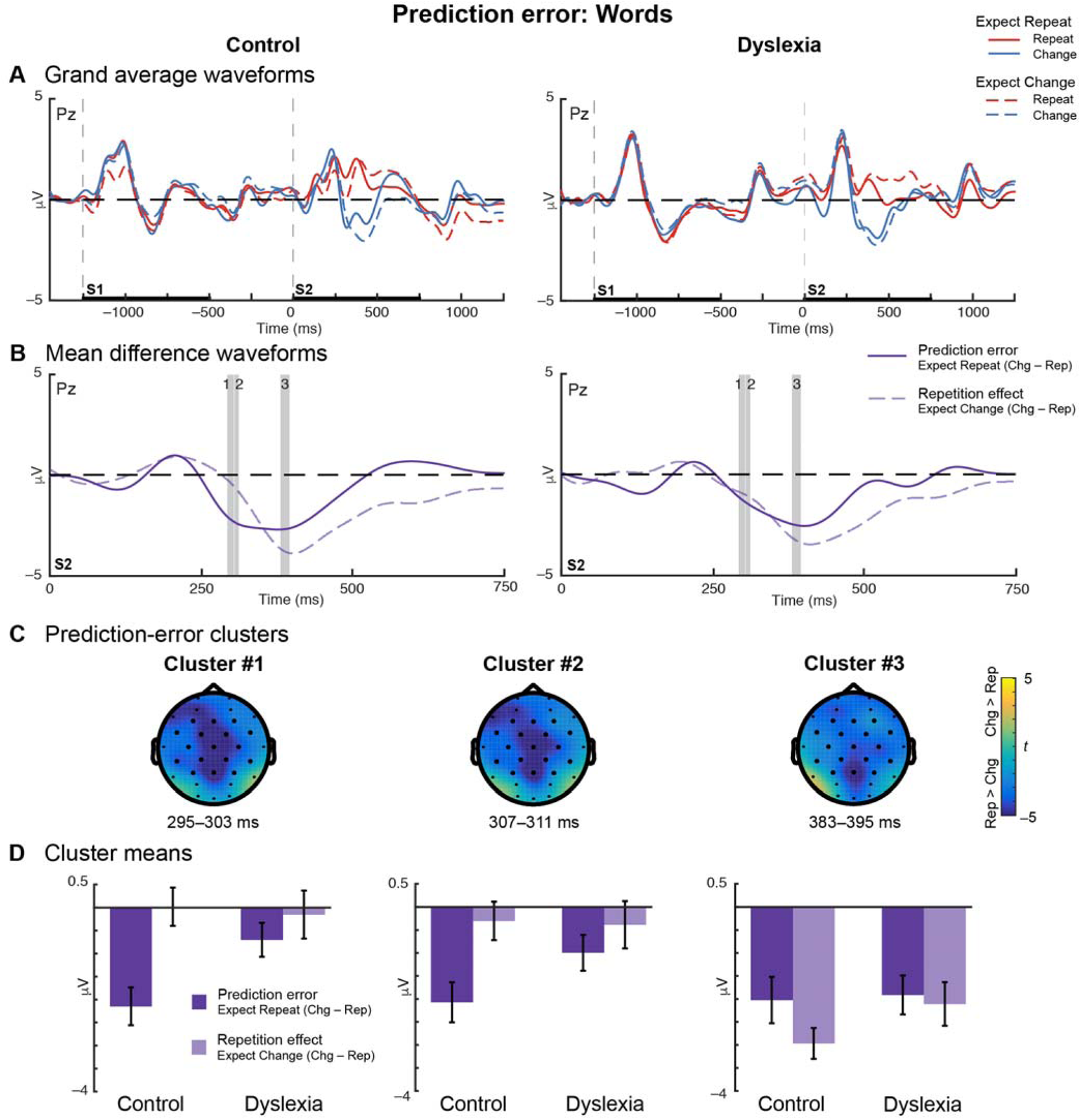
Reduced prediction error in dyslexia for unexpected changes versus expected repetitions of words. **(A)** Grand-average waveforms for Control (left) and Dyslexia (right) groups plotted at representative electrode Pz show that ERPs diverge during the second stimulus (S2) interval for Repetition (red) and Change (blue) trials under the expectation of repetition (solid lines) or of change (dashed lines). **(B)** Mean-difference waveforms for prediction error in the Expect Repeat condition (solid dark purple) and repetition in the Expect Change condition (dashed light purple) during S2 presentation. Control data are plotted on the left and Dyslexia on the right; gray bars on both indicate the durations of the prediction-error clusters identified in the Control group. **(C)** Topographical plots for each of the three prediction-error clusters identified in the Control group. Color indicates the prediction-error effect expressed as a *t*-statistic, averaged over the duration of the cluster. Dark electrodes significantly differentiate Change versus Repetition trials. **(D)** Mean-difference voltage values extracted from each cluster for Control and Dyslexia groups. Error bars represent (between-subjects) SEM. Prediction-error effects are accompanied by substantial repetition effects in Cluster #3. In Clusters #1 and 2, prediction error is significantly or trends larger in Control versus Dyslexia.

**Table 6.** Prediction-error effects (Words)

| Cluster | Time (ms) | Distribution | Polarity | ANOVA Results | $F_{1,38}$ | $p$ | |
| --- | --- | --- | --- | --- | --- | --- | --- |
| #1 | 295–303 | central | Rep > Chg | *Group* | 2.04 | 0.2 |  |
|  |  |  |  | *Repetition* | 8.87 | 0.005 | ** |
|  |  |  |  | *Expectation* | 1.05 | 0.3 |  |
|  |  |  |  | *Group × Repetition* | 1.62 | 0.2 |  |
|  |  |  |  | *Group × Expectation* | 0.48 | 0.5 |  |
|  |  |  |  | *Repetition × Expectation* | 15.01 | 0.0004 | *** |
|  |  |  |  | *Group × Repetition × Expectation* | 5.34 | 0.03 | * |
| #2 | 307–311 | central | Rep > Chg | *Group* | 1.98 | 0.2 |  |
|  |  |  |  | *Repetition* | 12.85 | 0.0009 | *** |
|  |  |  |  | *Expectation* | 1.48 | 0.2 |  |
|  |  |  |  | *Group × Repetition* | 0.89 | 0.4 |  |
|  |  |  |  | *Group × Expectation* | 1.19 | 0.3 |  |
|  |  |  |  | *Repetition × Expectation* | 12.45 | 0.001 | ** |
|  |  |  |  | *Group × Repetition × Expectation* | 2.98 | 0.09 |  |
| #3 | 383–395 | central | Rep > Chg | *Group* | 0.01 | 0.9 |  |
|  |  |  |  | *Repetition* | 85.41 | < 0.0001 | *** |
|  |  |  |  | *Expectation* | 16.77 | 0.0002 | *** |
|  |  |  |  | *Group × Repetition* | 0.99 | 0.3 |  |
|  |  |  |  | *Group × Expectation* | 1.74 | 0.2 |  |
|  |  |  |  | *Repetition × Expectation* | 2.22 | 0.1 |  |
|  |  |  |  | *Group × Repetition × Expectation* | 0.95 | 0.3 |  |

Clusters are numbered chronologically. Cluster time ranges are given with respect to the onset of S2. **Abbreviations:** Chg: Change trials; Rep: Repetition trials.

##### 3.4.2.1 Words Cluster #1

Repeated-measures ANOVA revealed a three-way *Group* × *Repetition* × *Expectation* interaction in the first cluster (**Table 6**). This interaction was driven by a robust prediction-error effect in the Control group (**Figure 5D, left**). Post-hoc tests revealed that the prediction-error effect was significantly larger in the Control group than in the Dyslexia group (*t* = −2.61; *p* = 0.01; *d* = 0.83), while the repetition effect did not differ significantly between groups (*t* = 0.26; *p* = 0.8; *d* = 0.08). In the Control group, the prediction-error effect was significantly larger than the repetition effect (*t* = −4.76; *p* = 0.0001; *d* = 1.17), but these two effects did not significantly differ in the Dyslexia group (*t* = −1.03; *p* = 0.3; *d* = 0.27). **Figure 5B** shows that the prediction-error effect for words has an earlier onset in the Control group, beginning ~300 ms after the onset of the unexpected word change.

##### 3.4.2.2 Words Cluster #2

The second cluster, which was nearly continuous with the first, showed a *Repetition* × *Expectation* interaction and a marginally-significant three-way interaction with *Group* (**Table 6**). A post-hoc test demonstrated that the magnitude of the prediction-error effect was larger than that of the repetition effect (*t* = −3.44; *p* = 0.001; *d* = 0.60) (**Figures 5B** and **5D, middle**)

##### 3.4.2.3 Words Cluster #3

The third cluster was characterized by main effects of *Repetition* and *Expectation* (**Table 6**; **Figure 5D, right**). The waveforms in **Figure 5B** demonstrate substantial and long-lasting prediction-error and repetition effects for word stimuli.

##### 3.4.2.4 Clusters defined in the Dyslexia group

When the ROI definition was performed in the Dyslexia group, Unexpected Changes and Expected Repetitions were found to be significantly different. Three broadly-distributed clusters were identified: 356–357, 383–414, and 420–432 ms (**Supp. Table 3**). Repeated-measures ANOVA on mean voltages extracted from the first cluster revealed main effects of *Repetition* and *Expectation*, while the second and third clusters both showed *Repetition* × *Expectation* interactions. Post-hoc tests indicated that the repetition effect was of a greater magnitude than the prediction-error effect in both the second (*t* = 2.13; *p* = 0.04; *d* = 0.40) and the third clusters (*t* = 2.72; *p* = 0.01; *d* = 0.47). There were no effects of nor interactions with *Group* in these three clusters (**Supp. Table 3**).

## 4. Discussion

The principal finding from this study is that neural prediction error, indexed by the difference in ERP magnitude between expected repetitions and unexpected changes, was diminished in adults with dyslexia compared to typically reading controls. This pattern of results was largely consistent across both face and text stimuli, suggesting that general-purpose, rather than domain-specific, cortical mechanisms for integrating top-down perceptual expectations with bottom-up sensory processing may be globally altered in dyslexia. We did not find evidence of a deficit in dyslexia for the feedforward, repetition-related neural response suppression that occurs after unexpected stimulus repetitions, suggesting that neural adaptation deficits in dyslexia may arise specifically due to a failure to integrate top-down expectation signals during perception, rather than dysfunction in bottom-up sensory processing. Finally, we did not find evidence of a domain-general deficit in dyslexia for generating top-down expectations about stimulus repetition, as indexed by modulation of prestimulus neural oscillatory activity, suggesting that the processes that generate top-down perceptual expectations are present in dyslexia.

### 4.1 Evidence against the Expectation-Deficit Hypothesis

Cortical oscillations reflect sustained, goal-directed attention to the environment, and fluctuations in the power of specific oscillations are related to variations in attention over time (Clayton, Yeung, & Kadosh, 2015). If individuals with dyslexia differed from typical readers in their capacity to allocate, control, or deploy top-down visual attention in expectation of different perceptual processing demands (Vidyasagar & Pammer, 2010; Facoetti et al., 2000), we would have expected to see differences between these groups in how prestimulus neural oscillatory activity is affected by anticipation of consistency vs. change in the perceptual environment (Vidyasagar, 2019). By creating two experimental conditions, one with a high probability of stimulus change and the other with a high probability of stimulus repetition, we effectively placed different attentional and information-processing demands on participants. In both participant groups, we observed that these differing demands significantly modulated activity in the theta and alpha frequency bands, concurrent with the presentation of the first stimulus in each pair and extending into the interstimulus interval. Relative to the Expect Change condition, the Expect Repeat condition was characterized by oscillatory desynchronization (i.e., a reduction in oscillatory power). Furthermore, the behavioral measures of response time and accuracy indicated that both groups were attentive to the task, irrespective of expectation condition or stimulus type.

For face stimuli, we found no significant differences in time-frequency representations between the Control and Dyslexia groups. This result suggests that the expectation of face repetition induces similar brain states in individuals with dyslexia compared to controls. For word stimuli, clusters of expectation-related neural desynchronization were also found in both groups. In one of the clusters defined in the Control group, the desynchronizing effect of expecting a word repetition was comparatively reduced in the Dyslexia group. Instead of difficulties generating perceptual expectations *per se*, this difference may reflect generally slower and less accurate word recognition in the Dyslexia group. The present task required rapid processing of the first stimulus to generate expectations about the upcoming one. It may be that, had longer processing time been available, we would have seen the magnitude of expectation-related neural states converge between the Control and Dyslexia groups. On the other hand, in one of the expectation-related time-frequency clusters defined in the Dyslexia group, we did not observe a corresponding desynchronizing effect in the Control group. This result suggests that generating perceptual expectations about text may rely on additional, distinct neural resources in dyslexia, which may reflect compensatory text-processing strategies not seen in controls (e.g., Hoeft et al., 2011).

The effects of neural oscillatory (de)synchronization on information processing are commonly studied via trial-by-trial memory performance. In contrast to the present study, in which participants were not required to encode the faces and words beyond monitoring for an inverted stimulus, many paradigms relate oscillatory power changes to the success with which a stimulus was remembered vs. forgotten (Sederberg et al., 2003; White et al., 2013). Prestimulus theta power enhancement has been associated with successful encoding of the events into memory, potentially by activating a memory context in which the stimulus can be encoded (Guderian et al., 2009; Fell et al., 2011; Gruber et al., 2013). Moreover, even infants demonstrate theta enhancement when they expect to receive information, e.g., from a speaker they can understand (Begus et al., 2016). These interpretations are consistent with the present results, in which enhanced theta power during the Expect Change condition relative to the Expect Repeat condition may reflect a neural state that is favorable for encoding new information vs. one where the demands for cognitive and perceptual resources are reduced, respectively.

Cortical oscillation in the alpha band has also been associated with attention, perception, and memory processes. Enhanced alpha oscillatory power is commonly considered to reflect the allocation of increased cognitive effort to a task through functional inhibition of the processing of task-irrelevant information or distractors (Klimesch, 2012; Jensen & Mazaheri, 2010; Snyder & Foxe, 2010; Strauss et al., 2014). Thus, alpha synchronization likely supports increasingly challenging visual tasks such as discrimination in the presence of distractors (Min & Herrmann, 2007) and retention of high working-memory loads (Jensen et al., 2002; Klimesch et al., 1999). Our finding of alpha modulation by expectation of a repeating vs. novel stimulus is consistent with this kind of differential demand on cognitive resources for perceptual processing: We observed relatively greater alpha power during the Expect Change condition (reflecting anticipation of the additional perceptual demands for processing a novel stimulus) and relatively reduced alpha power during the Expect Repeat condition (reflecting anticipation of the reduced perceptual demands for processing a repeated stimulus).

A caveat about this interpretation comes from the difference between the present task – in which participants had to detect rare deviant stimuli (inverted faces or words) – and the tasks used to study alpha (de)synchronization during differing attention and memory demands – in which participants have to suppress contemporaneous distractor stimuli. An important role for future work investigating putative visual-spatial attentional deficits in dyslexia will be to explore differences in alpha and theta neural oscillations in tasks analogous to those in the visual attention literature (e.g., Van der Lubbe, de Kleine, & Rataj, 2019). However, it is worth emphasizing that assessing visual-spatial attentional differences in dyslexia was not the aim of the present study. Instead, we sought to understand whether differences in the neural signatures of prestimulus attention and expectation would be different in dyslexia in a task where stimulus expectation should enhance neural adaptation (Summerfield et al., 2008; 2011), thus accounting for why neural adaptation to predictable stimulus repetition is diminished in dyslexia (Perrachione et al., 2016; Peter et al., 2019). Correspondingly, we found that individuals with dyslexia showed a fundamentally similar pattern of expectation-related neural oscillatory activity to controls: relative desynchronization under the condition in which repeated versus novel information was expected. From this we infer that the block-level manipulation of repetition probability was effective at inducing comparable top-down expectational states in both groups. This suggests that prior reports of differences in domain-general perceptual and neural adaptation in dyslexia do not merely reflect differences in the ability to develop top-down expectations about stimulus repetition, which have been shown to be critical for increasing repetition suppression magnitude (Larsson & Smith, 2012). Finally, it is also worth considering that, while expectation-related brain states measured by changes to EEG spectral power were largely similar between Control and Dyslexia groups, expectation-related brain states can be assessed with other types of signals, such as differences in neurotransmitter concentrations measured via pharmacological imaging (Bunzeck & Thiel, 2016), or activation of the locus-coeruleus system measured via pupillometry (Zhang et al., 2019), and future work may reveal group differences in expectation arising from other mechanisms.

### 4.2 Evidence against the Feedforward-Deficit Hypothesis

A classic finding across multiple methods of recording neural activity – from BOLD fMRI, to scalp electrophysiology, to recordings from individual neurons – is that repeated presentation of the same stimulus attenuates neural response (Grill-Spector et al., 2006). While the signal differences measured as population-level neural activity via neuroimaging doubtlessly reflect the aggregate change in response over many different mechanisms of short-term plasticity (Krekelberg et al., 2006; Larsson et al., 2016), some of these changes are strictly feedforward, in that they alter neural responses in the absence of top-down behavioral demands or when stimulus repetition is unexpected (e.g., Larsson & Smith, 2012). Instead of a failure to generate the top-down neuromodulatory signals that tune neuronal responses in expectation of particular stimulus features (as discussed above), the neural adaptation deficits previously observed in dyslexia could have been attributable to differences in strictly bottom-up processing that reduce the ability of population-level recordings like EEG and fMRI to detect repetition-induced changes in neural response. One prominent hypothesis is that feedforward neural responses are more stochastic in dyslexia (Hancock, Pugh, & Hoeft, 2017). These noisy, variable response profiles would lead to heterogeneous neural responses to the same stimulus across time (Hornickel & Kraus, 2013), weaker short-term representations of perceptual information (Jaffe-Dax et al., 2017; 2018), or both (Jaffe-Dax et al., 2015). Because stochastic neural responses activate slightly different neural populations each time a stimulus is encountered, there would appear to be less of an adaptation effect when aggregate neural responses are measured over large, undifferentiated neural populations of neurons via EEG or fMRI.

We evaluated this hypothesis by examining feedforward repetition effects in the absence of perceptual expectations, operationalized as the difference in response magnitude between Expected Change trials and Unexpected Repetition trials. That is, when participants do not have a top-down expectation of stimulus repetition, any difference between groups in the reduction of response magnitude between these two trial types can be attributed to differences in feedforward repetition suppression. It is worth considering that, in our paradigm, such unexpected repetitions were infrequent (25% of trials in that condition), but not so rare that participants were unaware they might happen. However, paradigms using similar rates of unexpected repetition have consistently shown that these events yield significantly smaller repetition-related response suppression compared to expected repetitions (e.g., Todorovic et al., 2011; Summerfield et al., 2008; 2011).

For face stimuli, we found no effect of feedforward repetition on ERP amplitude, neither in the Control group nor the Dyslexia group. That is, the response magnitude of an unexpected face repetition did not differ from an expected face change. While some studies have found repetition suppression for unexpected face repetitions (Summerfield et al., 2008), others have not – particularly when those faces were unfamiliar (Henson et al., 2002; Vizioli et al., 2010; reviewed in Schweinberger & Neumann, 2016). Given the diverging results from the prior literature, the present data may support the view that unfamiliar faces are not ideal stimuli to elicit a feedforward repetition-suppression response; however, given that expected repetitions of face stimuli did alter the magnitude of evoked responses (discussed below), it may instead be that the present study was underpowered to detect feedforward repetition effects for unfamiliar faces.

In contrast, we found long-lasting and robust feedforward repetition effects in the ERP signal for word stimuli, beginning at 365 ms after stimulus onset. These effects were identified separately in both the Control and Dyslexia groups. Of the seven spatiotemporal clusters identified across the two groups, six showed no statistical difference between groups, and one showed an effect of repetition in the Dyslexia group alone. These results demonstrate that, in adults with dyslexia, an unpredicted second exposure to a short, written word is sufficient to induce a neural repetition effect quantitatively similar to that measured in typical adult readers.

The lack of a group difference in feedforward repetition suppression provides an important clarifying perspective on previous literature showing that individuals with dyslexia respond differently to stimulus consistency, whether measured in their behavior (Ahissar et al., 2006) or brain responses (Baldeweg et al., 1999; Stoodley et al., 2006; Perrachione et al., 2016; Chandrasekaran et al., 2009). Our ERP findings suggest that, in dyslexia, feedforward sensory processing remains sensitive to repetition. Intact feedforward adaptation responses pose a challenge for theories of dyslexia that posit greater stochasticity in the feedforward neural responses to a consistent sensory input (Hancock et al., 2017), because many of the mechanisms proposed to underlie feedforward repetition suppression depend on consistent reactivation of the same neural populations (Kohn & Movshon, 2003; Kohn & Movshon, 2004; Kaliukhovich & Vogels, 2010; reviewed in Vogels, 2016). However, a “neural noise” hypothesis applies equally to stochasticity of feed*back* responses as it does to feedforward ones. As such, the reduction in adaptation may be the result poor or noisy timing for the integration of top-down signals conveying an expectation of repetition with feedforward signals conveying repeated stimulus features – a possibility we consider below.

### 4.3 Evidence for the Expectation Integration-Deficit Hypothesis

Generating top-down expectations of perceptual experiences is of little use in facilitating perceptual processing if these signals are not successfully integrated with bottom-up sensory representations. Incorporating perceptual expectations into feedforward sensory processing serves two important purposes: First, via predictive coding, it reduces the enormous physiological cost of continuously processing an environment filled with static signals that have little relevance for behavior; second, via prediction error, it provides a mechanism for learning when perceptual expectations are violated (Press et al., 2020; Grotheer & Kovacs, 2016; Auksztulewicz & Friston, 2016). In particular, expectations about perceptual events tend to lead to suppression of neural activity (also known as *expectation suppression*; Todorovic et al., 2011), which was first noted as a critical process in sensorimotor integration and motor learning, as organisms must be able to dissociate sensory experiences related to their own actions from those that arise externally from the environment (Crapse & Sommer, 2008). A mismatch between expectations and sensory experiences generates an error signal that not only reorients attention to relevant external stimuli, but also provides a mechanism for neural plasticity as an organism learns to make better predictions. For example, prominent models of speech motor learning are based on such perceptual error-driven plasticity mechanisms (Guenther, 2016).

In this study, we leveraged the relationship between expectation suppression and prediction error to investigate whether there is an expectation-integration deficit in dyslexia. By comparing the difference in neural evoked response to stimuli that fulfilled perceptual expectations (Expected Repetition trials) versus stimuli that violated those expectations (Unexpected Change trials), we were able to investigate how perceptual predictive coding differentially affected neural response dynamics between groups. Specifically, we identified spatiotemporal clusters in the Control group where neural responses reflected prediction error, then investigated whether the response profiles within these clusters differed in the Dyslexia group as a function of repetition expectation.

For face stimuli, we found three clusters with prediction-error effects in Controls. Neural responses to repetition in the first cluster (258–305ms) were also significantly affected by expectation (with stronger prediction-error than repetition effects) and by group (with larger effects in Controls than in Dyslexia). In the second cluster (391–406ms), the repetition effect was only marginally affected by expectation or by group. However, in the third cluster (428–440ms), not only was the repetition effect significantly modulated by expectation (with stronger prediction-error responses), there was also a significant three-way group by repetition by expectation interaction, such that expectation of stimulus repetition had a smaller effect on the repetition-related change in neural response in Dyslexia than in the Controls. (As seen in **Figures 4B** and **4C**, the topographic distributions of the latter two clusters are virtually identical, and they are highly proximate in time. However, we chose to analyze these as distinct clusters in keeping with the standards of data-driven analysis.) Similarly, for word stimuli, we identified three spatiotemporal clusters with significant prediction-error effects in the Control group. In the first of these clusters (295–303ms), not only was the repetition effect modulated by expectation, the degree of this modulation was different between groups: Expectation of stimulus repetition had a considerably smaller effect on neural responses to repetition in individuals with dyslexia compared to controls. The second cluster (307–311ms) was temporally proximal and topographically similar to the preceding cluster. As before, we chose to analyze these separately in keeping with our data-driven methods; however, mechanistically, we believe the response profiles in these clusters likely reflect similar neural populations. In this cluster, the effect of repetition on neural response was again significantly moderated by participants’ expectations, though the difference in this effect across groups was smaller than in the preceding cluster. (In a third cluster (383–395ms), there was no indication of difference between the two groups.) In sum, for both face and word stimuli, neural responses to repetition are modulated differently by perceptual expectations between individuals with and without dyslexia.

These findings suggest that individuals with dyslexia have a specific weakness in integrating top-down expectations about future stimuli during perceptual processing. When participants could not predict the upcoming stimulus repetition, we observed robust bottom-up repetition suppression in both groups. The magnitude and timecourse of neural responses to unexpected repetition effects in both groups were similarly weak for faces and robust for words. In contrast, when a prediction was available, we observed significantly weaker prediction error in the Dyslexia group. Intuitively, the effects of repetition, expectation, and group can be appreciated from the difference waves in **Figures 4B** (faces) and **5B** (words), in which each trace depicts the ERP difference between Change trials and Repetition trials. At certain times, the magnitude of that difference is exaggerated in the Expect Repeat condition relative to the Expect Change condition, consistent with the heuristic that Unexpected Changes yield higher highs and Expected Repetitions yield lower lows of population-level neural response.

It may be the case that, in dyslexia, top-down prediction signals are less effective at tuning feedforward sensory processing, reducing perceptual efficiency and posing additional neurocomputational, and thus physiological, costs on perception. The effective communication of top-down expectations may be disrupted by structural or functional disconnection between the higher-order cortical areas that generate those signals and the lower-order cortical areas that integrate them with feedforward sensory representations (Boets et al., 2013; Saygin et al., 2013; Yeatman et al., 2011). Alternately, local disorganization of the cortical microstructure in dyslexia may affect the efficacy with which top-down signals can tune feedforward responses (Galaburda et al., 1994; 2006). Ongoing research both with animal models and human neuroimaging is beginning to reveal how specific neurotransmitter systems are responsible for tuning neural responses in sensory cortices on short timescales based on top-down neuromodulatory inputs, which may provide foundations for future inquiries into the neurochemical foundations of learning difficulties in dyslexia (Froemke et al., 2007; Fritz et al., 2003; Bunzeck & Thiel, 2016).

The present results also offer new insight into our prior observations of widespread neural adaptation deficits in individuals with dyslexia. Previously, using fMRI, we found evidence of reduced neural adaptation to faces and written words, among many types of stimuli (Perrachione et al., 2016). Specifically, in adults and children with dyslexia, there was less of a difference in the magnitude of the BOLD response between blocks where one stimulus was repeated multiple times in a row vs. single presentations of many different stimuli. These conditions are analogous to the “Expect Repeat” and “Expect Change” conditions of the present study, as participants could quickly and accurately generate valid predictions for whether upcoming stimuli would be repetitions vs. novel. Prior work has shown that the magnitude of BOLD adaptation is affected by expectations about stimulus repetition, with unexpected repetitions leading to a smaller reduction in BOLD response compared to expected repetitions (Summerfield et al., 2008). Similarly, although its magnitude is less than in controls, low levels of neural adaptation are still seen in individuals with dyslexia (Perrachione et al., 2016), consistent with our observations in the present study of intact bottom-up repetition suppression in both groups, but a reduced effect of perceptual expectation on repetition-related responses in dyslexia. Thus, in the prior work, it may be that reduced neural adaptation in dyslexia measured via fMRI reflects the failure to successfully integrate (conscious) top-down expectations about the likelihood of stimulus repetition with the bottom-up sensory processes responsible for encoding perceptual representations.

These results may also help clarify adaptation-like deficits in electrophysiological studies of dyslexia, particularly those measuring the MMN component. To measure the MMN response, a long series of adapting stimuli is typically presented prior to an unpredictable deviant stimulus. Individuals with dyslexia have consistently been found to have smaller MMN responses to a rare deviant compared to typical readers (Gu & Bi, 2020; Hämäläinen, Salminen, & Leppänen, 2012; Schulte-Körne & Bruder, 2010). However, prior MMN studies in dyslexia have not attempted to ascertain to what extent smaller MMN responses in dyslexia are attributable to weaker bottom-up versus top-down effects. Furthermore, the design of MMN-eliciting paradigms makes it difficult to disentangle bottom-up and top-down effects, and competing computational and neurobiological accounts of this phenomenon have variously emphasized its causal origins as a short-term memory trace (Näätänen, Jacobsen, & Winkler, 2005), an adaptation process (May & Tiitinen, 2010), or a representation of the environment’s statistical structure (Herrmann et al., 2015). By and large, each of these explanations has a parallel in a model based on predictive coding and prediction error (Baldeweg, 2007), which provides a parsimonious framework for understanding both the MMN and, informed by the present results, its reduction in dyslexia. It is worth noting that atypical mismatch responses are also found in newborns with a family history of dyslexia (Leppänen et al., 1999), further suggesting that circuit-level differences may have genetic, rather than experiential, origins in dyslexia.

### 4.4 Expectation integration, prediction error, and learning to read in dyslexia

Reduced capacity for expectation integration and the consequent attenuation of prediction error have theoretical importance for how we understand various dyslexia phenotypes and their etiologies. A first major consequence of an expectation-integration deficit is reduced neural efficiency. Stimulus repetition facilitates perception (Maccotta & Buckner, 2004) – but so too does repeated presentation of various partial views of a stimulus (Doniger et al., 2001), suggesting not only that a complete, high-level representation is built through evidence accumulation, but that such a representation feeds backward and expedites object recognition. By a similar principle, semantic primes (e.g., a word and a picture) induce neural adaptation for one another in ventral occipitotemporal cortex even though they do not share low-level features (Kherif, Josse, & Price, 2010). Reduced neural efficiency due to impaired prediction may explain several of the phenotypes of dyslexia. For example, analysis of eye movements reveals that fast readers make more predictions, or “forward inferences” than do slow readers (Hawelka et al., 2015). Moreover, the prediction may not be limited to the visual/orthographic features of the upcoming word, but may also include its semantic and phonological features – these three together comprising a high-quality word representation that supports reading comprehension (Perfetti, 2007). This pattern may extend to auditory language processing as well, with evidence that individuals with dyslexia are slower to direct their gaze to targets cued by the grammar (e.g., gender marker) of spoken instructions (Huettig & Brouwer, 2015). General weaknesses in the ability to predict “what comes next” have been widely documented in dyslexia, including on the serial reaction time task (an implicit, sequenced motor skill: Lum et al., 2013) and in a first-person shooter video game in which implicitly-learned auditory categories probabilistically cue target appearance (Gabay & Holt, 2015). There is evidence that successful reading acquisition is associated with the ability to take advantage of successful predictions to improve reading fluency, such as fewer eye fixations and fewer regressive eye movements (Starr & Rayner, 2001), and that typically reading children take advantage of statistical regularities in text to improve accuracy and fluency more so than readers with dyslexia (Jones et al., 2020; Franzen et al., 2021).

Prediction is useful not only because it pre-activates perceptual information, but also because it serves as a template against which to compare incoming signals: Consistent sensory information is processed more efficiently, while a mismatch triggers an error response that drives plasticity (Press et al., 2020). Price and Devlin (2011) describe how the magnitude of prediction error varies during learning: Initially, no learning occurs because a novel, meaningless stimulus has no existing representation, generates no prediction, and yields no prediction error. During learning, the stimulus becomes familiar but is not efficiently predicted, and large prediction errors serve to build its representation. Finally, with expertise, representations are robust, predictions are generally accurate, and small prediction errors simply refine those predictions in new contexts. Similar trajectories are seen during reading acquisition, as children learn to balance recognition, accuracy, and fluency in the decoding of text. Such learning, however, will be disproportionately challenged if the mechanism for integrating predictions into feedforward processing is faulty, leading to less reliable or less effective generation of prediction errors. Without prediction error, learning signals cannot be sent onwards to trigger plasticity, refine predictions, and ultimately build the long-term representations and statistical associations that underlie complex perceptual tasks like accurate and fluent reading. In other words, the second major consequence of an expectation-integration deficit is reduced learning. Multiple lines of evidence, including the present results, now suggest that the representation of statistical regularities, and recognizing deviations from them, may be impaired in dyslexia (Menghini et al., 2006; Stoodley et al., 2008; Sperling et al., 2005; Jones et al., 2020) and may play a role in these individuals’ difficulty attaining accurate and fluent reading skills.

## 5. Conclusions

In dyslexia, stimulus repetition has been shown to result in less behavioral facilitation and less neural adaptation compared to typical readers. Here, we showed that diminished repetition-related neural responses in dyslexia may be specifically related to a failure to integrate top-down expectations of stimulus repetition with bottom-up sensory encoding, rather than differences in top-down expectation or bottom-up encoding themselves. Attenuation of the neural correlate of perceptual prediction error in dyslexia is a candidate for the sort of subtle dysfunction in neural systems for learning that may impede the development of accurate and fluent reading skills.

## 6. Acknowledgments

We thank Rebecca Winter, Stephanie Del Tufo, Bianca Levy, and Patricia Chang for their assistance with data collection and Steve Shannon and Sheeba Arnold at the Athinoula A. Martinos Imaging Center at the McGovern Institute for Brain Research at MIT for technical support. Research reported in this article was supported by the Ellison Medical Foundation and the National Institutes of Health grants UL1RR025758 (to JDEG) and R03HD096098 (to TKP). SDB was supported by NIH grants T32DC000038 and F31HD100101 and by the Friends of the McGovern Institute Fellowship. SJL was supported by NIH grant T32DC013017. The content is solely the responsibility of the authors and does not necessarily represent the official views of the National Institutes of Health.

## Footnotes

1 Example stimuli may be found at http://www.emotionlab.se/resources/kdef (Karolinska), http://www.macbrain.org/resources.htm (NimStim), and http://www.socsci.ru.nl:8180/RaFD2/RaFD (Radboud).

## References

Ahissar, M., Lubin, Y., Putter-Katz, H., & Banai, K. (2006). Dyslexia and the failure to form a perceptual anchor. Nature Neuroscience, 9(12), 1558.

Arciuli, J., & Simpson, I. C. (2012). Statistical learning is related to reading ability in children and adults. Cognitive Science, 36(2), 286–304.

Atiani, S., Elhilali, M., David, S. V., Fritz, J. B., & Shamma, S. A. (2009). Task difficulty and performance induce diverse adaptive patterns in gain and shape of primary auditory cortical receptive fields. Neuron, 61(3), 467–480.

Auksztulewicz, R., & Friston, K. (2016). Repetition suppression and its contextual determinants in predictive coding. Cortex, 80, 125–140.

Baldeweg, T. (2007). ERP repetition effects and mismatch negativity generation: A predictive coding perspective. Journal of Psychophysiology, 21(3-4), 204–213.

Baldeweg, T., Richardson, A., Watkins, S., Foale, C., & Gruzelier, J. (1999). Impaired auditory frequency discrimination in dyslexia detected with mismatch evoked potentials. Annals of Neurology, 45(4), 495–503.

Begus, K., Gliga, T., & Southgate, V. (2016). Infants’ preferences for native speakers are associated with an expectation of information. Proceedings of the National Academy of Sciences, 113(44), 12397–12402.

Boets, B., de Beeck, H. P. O., Vandermosten, M., Scott, S. K., Gillebert, C. R., Mantini, D., … & Ghesquière, P. (2013). Intact but less accessible phonetic representations in adults with dyslexia. Science, 342(6163), 1251–1254.

Brunet, N. M., Bosman, C. A., Vinck, M., Roberts, M., Oostenveld, R., Desimone, R., De Weerd, P., & Fries, P. (2014). Stimulus repetition modulates gamma-band synchronization in primate visual cortex. Proceedings of the National Academy of Sciences, 111(9), 3626–3631.

Bunzeck, N., & Thiel, C. (2016). Neurochemical modulation of repetition suppression and novelty signals in the human brain. Cortex, 80, 161–173.

Centanni, T. M., Pantazis, D., Truong, D. T., Gruen, J. R., Gabrieli, J. D. E., & Hogan, T. P. (2018). Increased variability of stimulus-driven cortical responses is associated with genetic variability in children with and without dyslexia. Developmental Cognitive Neuroscience, 34, 7–17.

Chait, M., Eden, G., Poeppel, D., Simon, J. Z., Hill, D. F., & Flowers, D. L. (2007). Delayed detection of tonal targets in background noise in dyslexia. Brain and Language, 102(1), 80–90.

Chandrasekaran, B., Hornickel, J., Skoe, E., Nicol, T., & Kraus, N. (2009). Context-dependent encoding in the human auditory brainstem relates to hearing speech in noise: Implications for developmental dyslexia. Neuron, 64(3), 311–319.

Clark, A. (2013). Whatever next? Predictive brains, situated agents, and the future of cognitive science. Behavioral and Brain Sciences, 36(3), 181–204.

Clayton, M. S., Yeung, N., & Kadosh, R. C. (2015). The roles of cortical oscillations in sustained attention. Trends in Cognitive Sciences, 19(4), 188–195.

Crapse, T. B. & Sommer, M. A. (2008). Corollary discharge across the animal kingdom. Nature Reviews Neuroscience, 9(8), 587–600.

Dehaene, S., & Cohen, L. (2007). Cultural recycling of cortical maps. Neuron, 56(2), 384–398.

Delorme, A., & Makeig, S. (2004). EEGLAB: an open source toolbox for analysis of single-trial EEG dynamics including independent component analysis. Journal of Neuroscience Methods, 134(1), 9–21.

Desimone, R. (1996). Neural mechanisms for visual memory and their role in attention. Proceedings of the National Academy of Sciences, 93(24), 13494–13499.

Doniger, G. M., Foxe, J. J., Schroeder, C. E., Murray, M. M., Higgins, B. A., & Javitt, D. C. (2001). Visual perceptual learning in human object recognition areas: A repetition priming study using high-density electrical mapping. Neuroimage, 13(2), 305–313.

Facoetti, A., Paganoni, P., Turatto, M., Marzola, V., & Mascetti, G. G. (2000). Visual-spatial attention in developmental dyslexia. Cortex, 36(1), 109–123.

Fell, J., Ludowig, E., Staresina, B. P., Wagner, T., Kranz, T., Elger, C. E., & Axmacher, N. (2011). Medial temporal theta/alpha power enhancement precedes successful memory encoding: Evidence based on intracranial EEG. Journal of Neuroscience, 31(14), 5392–5397.

Franzen, L., Stark, Z., & Johnson, A. P. (2021). Individuals with dyslexia use a different visual sampling strategy to read text. Scientific Reports, 11(1), 1–17.

Friston, K. (2009). The free-energy principle: A rough guide to the brain? Trends in Cognitive Sciences, 13(7), 293–301.

Fritz, J., Shamma, S., Elhilali, M., & Klein, D. (2003). Rapid task-related plasticity of spectrotemporal receptive fields in primary auditory cortex. Nature Neuroscience, 6(11), 1216–1223.

Froemke, R. C., Merzenich, M. M., & Schreiner, C. E. (2007). A synaptic memory trace for cortical receptive field plasticity. Nature, 450(7168), 425–429.

Gabay, Y., & Holt, L. L. (2015). Incidental learning of sound categories is impaired in developmental dyslexia. Cortex, 73, 131–143.

Gabay, Y., & Holt, L. L. (2020). Adaptive plasticity under adverse listening conditions in developmental dyslexia. Journal of the International Neuropsychological Society, 1–11.

Gabrieli, J.D.E. (2009) Dyslexia: A new synergy between education and cognitive neuroscience. Science, 325, 280–283.

Galaburda, A.M., LoTurco, J., Ramus, F., Fitch, R.H., & Rosen, G.D. (2006). From genes to behavior in developmental dyslexia. Nature Neuroscience, 9, 1213–1217.

Galaburda, A.M., Menard, M.T., & Rosen, G.D. (1994). Evidence for aberrant auditory anatomy in developmental dyslexia. Proceedings of the National Academy of Sciences, 91, 8010–8013.

Garrido, M. I., Kilner, J. M., Stephan, K. E., & Friston, K. J. (2009). The mismatch negativity: A review of underlying mechanisms. Clinical Neurophysiology, 120(3), 453–463.

Grill-Spector, K., Henson, R., & Martin, A. (2006). Repetition and the brain: Neural models of stimulus-specific effects. Trends in Cognitive Sciences, 10(1), 14–23.

Grotheer, M., & Kovács, G. (2016). Can predictive coding explain repetition suppression? Cortex, 80, 113–124.

Gruber, M. J., Watrous, A. J., Ekstrom, A. D., Ranganath, C., & Otten, L. J. (2013). Expected reward modulates encoding-related theta activity before an event. NeuroImage, 64(6), 68–74.

Gu, C. & Bi, H.-Y. (2020). Auditory processing deficit in individuals with dyslexia: A meta-analysis of mismatch negativity. Neuroscience & Biobehavioral Reviews, 116, 396–405.

Guderian, S., Schott, B. H., Richardson-Klavehn, A., & Düzel, E. (2009). Medial temporal theta state before an event predicts episodic encoding success in humans. Proceedings of the National Academy of Sciences, 106(13), 5365–5370.

Guenther, F. H. (2016). Neural Control of Speech. Cambridge, MA: MIT Press.

Hämäläinen, J. A., Salminen, H. K., & Leppänen, P. H. (2013). Basic auditory processing deficits in dyslexia: systematic review of the behavioral and event-related potential/field evidence. Journal of Learning Disabilities, 46(5), 413–427.

Hancock, R., Pugh, K. R., & Hoeft, F. (2017). Neural noise hypothesis of developmental dyslexia. Trends in Cognitive Sciences, 21(6), 434–448.

Hansen, B. J., & Dragoi, V. (2011). Adaptation-induced synchronization in laminar cortical circuits. Proceedings of the National Academy of Sciences, 108(26), 10720–10725.

Harm, M. W., & Seidenberg, M. S. (2004). Computing the meanings of words in reading: Cooperative division of labor between visual and phonological processes. Psychological Review, 111(3), 662.

Hawelka, S., Schuster, S., Gagl, B., & Hutzler, F. (2015). On forward inferences of fast and slow readers. An eye movement study. Scientific Reports, 5, 8432.

Henson, R. N. A. (2003). Neuroimaging studies of priming. Progress in Neurobiology 70, 53–81.

Henson, R. N. A., Shallice, T., Gorno-Tempini, M. L., & Dolan, R. J. (2002). Face repetition effects in implicit and explicit memory tests as measured by fMRI. Cerebral Cortex, 12(2), 178–186.

Herrmann, B., Henry, M. J., Fromboluti, E. K., McAuley, J. D., & Obleser, J. (2015). Statistical context shapes stimulus-specific adaptation in human auditory cortex. Journal of Neurophysiology, 113(7), 2582–2591.

Hoeft, F., McCandliss, B. D., Black, J. M., Gantman, A., Zakerani, N., Hulme, C., Lyytinen, H., Whitfield-Gabrieli, S., Glover, G. H., Reiss, A. L. Gabrieli, J. D. E. (2011). Neural systems predicting long-term outcome in dyslexia. Proceedings of the National Academy of Sciences, 108(1), 361–366.

Hornickel, J., & Kraus, N. (2013). Unstable representation of sound: A biological marker of dyslexia. Journal of Neuroscience, 33(8), 3500–3504.

Huettig, F., & Brouwer, S. (2015). Delayed anticipatory spoken language processing in adults with dyslexia—evidence from eyeutracking. Dyslexia, 21(2), 97–122.

Jaffe-Dax, S., Kimel, E., & Ahissar, M. (2018). Shorter cortical adaptation in dyslexia is broadly distributed in the superior temporal lobe and includes the primary auditory cortex. eLife, 7, e30018.

Jaffe-Dax, S., Lieder, I., Biron, T., & Ahissar, M. (2016). Dyslexics’ usage of visual priors is impaired. Journal of Vision, 16(9), 10–10.

Jaffe-Dax, S., Frenkel, O., & Ahissar, M. (2017). Dyslexics’ faster decay of implicit memory for sounds and words is manifested in their shorter neural adaptation. eLife, 6, e20557.

Jaffe-Dax, S., Raviv, O., Jacoby, N., Loewenstein, Y., & Ahissar, M. (2015). A computational model of implicit memory captures dyslexics’ perceptual deficits. Journal of Neuroscience, 35(35), 12116–12126.

Jensen, O., & Mazaheri, A. (2010). Shaping functional architecture by oscillatory alpha activity: Gating by inhibition. Frontiers in Human Neuroscience, 4, 186.

Jensen, O., Gelfand, J., Kounios, J., & Lisman, J. E. (2002). Oscillations in the alpha band (9–12 Hz) increase with memory load during retention in a short-term memory task. Cerebral Cortex, 12(8), 877–882.

Jones, M., Kuipers, J. R., Nugent, S., Miley, A., & Oppenheim, G. (2018). Episodic traces and statistical regularities: Paired associate learning in typical and dyslexic readers. Cognition, 177, 214–225.

Kaliukhovich, D. A., & Vogels, R. (2010). Stimulus repetition probability does not affect repetition suppression in macaque inferior temporal cortex. Cerebral Cortex, 21(7), 1547–1558.

Kanwisher, N., McDermott, J., & Chun, M. M. (1997). The fusiform face area: A module in human extrastriate cortex specialized for face perception. Journal of Neuroscience, 17(11), 4302–4311.

Kastner, S., Pinsk, M. A., De Weerd, P., Desimone, R., & Ungerleider, L. G. (1999). Increased activity in human visual cortex during directed attention in the absence of visual stimulation. Neuron, 22(4), 751–761.

Khalighinejad, B., Herrero, J. L., Mehta, A. D., & Mesgarani, N. (2019). Adaptation of the human auditory cortex to changing background noise. Nature Communications, 10(1), 2509.

Kherif, F., Josse, G., & Price, C. J. (2010). Automatic top-down processing explains common left occipito-temporal responses to visual words and objects. Cerebral Cortex, 21(1), 103–114.

Klimesch, W. (2012). Alpha-band oscillations, attention, and controlled access to stored information. Trends in Cognitive Sciences, 16(12), 606–617.

Klimesch, W., Doppelmayr, M., Schwaiger, J., Auinger, P., & Winkler, T. (1999). Paradoxical alpha synchronization in a memory task. Cognitive Brain Research, 7(4), 493–501.

Kohn, A., & Movshon, J. A. (2003). Neuronal adaptation to visual motion in area MT of the macaque. Neuron, 39(4), 681–691.

Kohn, A., & Movshon, J. A. (2004). Adaptation changes the direction tuning of macaque MT neurons. Nature Neuroscience, 7(7), 764.

Kok, P., Failing, M. F., & de Lange, F. P. (2014). Prior expectations evoke stimulus templates in the primary visual cortex. Journal of Cognitive Neuroscience, 26(7), 1546–1554.

Krekelberg, B., Boynton, G. M., & van Wezel, R. J. (2006). Adaptation: From single cells to BOLD signals. Trends in Neurosciences, 29(5), 250–256.

Larsson, J., & Smith, A. T. (2012). fMRI repetition suppression: Neuronal adaptation or stimulus expectation? Cerebral Cortex, 22(3), 567–576.

Larsson, J., Solomon, S. G., & Kohn, A. (2016). fMRI adaptation revisited. Cortex, 80, 154–160.

Leppänen, P. H., Pihko, E., Eklund, K. M., & Lyytinen, H. (1999). Cortical responses of infants with and without a genetic risk for dyslexia: II. Group effects. Neuroreport, 10(5), 969–973.

Linkersdörfer, J., Lonnemann, J., Lindberg, S., Hasselhorn, M., & Fiebach, C. J. (2012). Grey matter alterations co-localize with functional abnormalities in developmental dyslexia: An ALE meta-analysis. PloS one, 7(8), e43122.

Lum, J. A., Ullman, M. T., & Conti-Ramsden, G. (2013). Procedural learning is impaired in dyslexia: Evidence from a meta-analysis of serial reaction time studies. Research in Developmental Disabilities, 34(10), 3460–3476.

Lyon, G. R., Shaywitz, S. E., & Shaywitz, B. A. (2003). A definition of dyslexia. Annals of Dyslexia, 53(1), 1–14.

Maccotta, L., & Buckner, R. L. (2004). Evidence for neural effects of repetition that directly correlate with behavioral priming. Journal of Cognitive Neuroscience, 16(9), 1625–1632.

Maris, E., & Oostenveld, R. (2007). Nonparametric statistical testing of EEG- and MEG-data. Journal of Neuroscience Methods, 164(1), 177–190.

Marlin, S. G., Hasan, S. J., Cynader, M. S. (1988). Direction-selective adaptation in simple and complex cells in cat striate cortex. Journal of Neurophysiology, 59(4), 1314–1330.

Martin, A., Kronbichler, M., & Richlan, F. (2016). Dyslexic brain activation abnormalities in deep and shallow orthographies: A metauanalysis of 28 functional neuroimaging studies. Human Brain Mapping, 37(7), 2676–2699.

Maurer, U., Bucher, K., Brem, S., & Brandeis, D. (2003). Altered responses to tone and phoneme mismatch in kindergartners at familial dyslexia risk. Neuroreport, 14(17), 2245–2250.

May, P. J., & Tiitinen, H. (2010). Mismatch negativity (MMN), the devianceuelicited auditory deflection, explained. Psychophysiology, 47(1), 66–122.

McCandliss, B. D., Cohen, L., & Dehaene, S. (2003). The visual word form area: Expertise for reading in the fusiform gyrus. Trends in Cognitive Sciences, 7(7), 293–299.

Menghini, D., Hagberg, G. E., Caltagirone, C., Petrosini, L., & Vicari, S. (2006). Implicit learning deficits in dyslexic adults: An fMRI study. NeuroImage, 33(4), 1218–1226.

Min, B. K., & Herrmann, C. S. (2007). Prestimulus EEG alpha activity reflects prestimulus top-down processing. Neuroscience Letters, 422(2), 131–135.

Näätänen, R., Jacobsen, T., & Winkler, I. (2005). Memory based or afferent processes in mismatch negativity (MMN): A review of the evidence. Psychophysiology, 42(1), 25–32.

Neuhoff, N., Bruder, J., Bartling, J., Warnke, A., Remschmidt, H., Müller-Myhsok, B., & Schulte-Körne, G. (2012). Evidence for the late MMN as a neurophysiological endophenotype for dyslexia. PloS one, 7(5), e34909.

Oganian, Y., & Ahissar, M. (2012). Poor anchoring limits dyslexics’ perceptual, memory, and reading skills. Neuropsychologia, 50(8), 1895–1905.

Oostenveld, R., Fries, P., Maris, E., & Schoffelen, J. M. (2011). FieldTrip: Open source software for advanced analysis of MEG, EEG, and invasive electrophysiological data. Computational Intelligence and Neuroscience, 2011, 1.

Perea, M., Jiménez, M., Suárez-Coalla, P., Fernández, N., Viña, C., & Cuetos, F. (2014). Ability for voice recognition is a marker for dyslexia in children. Experimental Psychology, 61, 480–487.

Perfetti, C. (2007). Reading ability: Lexical quality to comprehension. Scientific Studies of Reading, 11(4), 357–383.

Perrachione, T. K., Del Tufo, S. N., & Gabrieli, J. D. (2011). Human voice recognition depends on language ability. Science, 333(6042), 595–595.

Perrachione, T. K., Del Tufo, S. N., Winter, R., Murtagh, J., Cyr, A., Chang, P., Halverson, K., Ghosh, S. S., Christodoulou, J. A., & Gabrieli, J. D. (2016). Dysfunction of rapid neural adaptation in dyslexia. Neuron, 92(6), 1383–1397.

Peter, B., McCollum, H., Daliri, A., & Panagiotides, H. (2019). Auditory gating in adults with dyslexia: An ERP account of diminished rapid neural adaptation. Clinical Neurophysiology, 130(11), 2182–2192.

Press, C., Kok, P., & Yon, D. (2020). The perceptual prediction paradox. Trends in Cognitive Sciences, 24(1), 13–24.

Price, C. J., & Devlin, J. T. (2011). The interactive account of ventral occipitotemporal contributions to reading. Trends in Cognitive Sciences, 15(6), 246–253.

Ramus, F. & Szenkovits, G. (2008). What phonological deficit? Quarterly Journal of Experimental Psychology, 61(1), 129–141.

Rao, R. P., & Ballard, D. H. (1999). Predictive coding in the visual cortex: A functional interpretation of some extra-classical receptive-field effects. Nature Neuroscience, 2(1), 79.

Saygin, Z. M., Norton, E. S., Osher, D. E., Beach, S. D., Cyr, A. B., Ozernov-Palchik, O., … & Gabrieli, J. D. (2013). Tracking the roots of reading ability: White matter volume and integrity correlate with phonological awareness in prereading and early-reading kindergarten children. Journal of Neuroscience, 33(33), 13251–13258.

Schacter, D. L., & Buckner, R. L. (1998). Priming and the brain. Neuron, 20(2), 185–195.

Schulte-Körne, G., & Bruder, J. (2010). Clinical neurophysiology of visual and auditory processing in dyslexia: A review. Clinical Neurophysiology, 121(11), 1794–1809.

Schulte-Körne, G., Deimel, W., Bartling, J., & Remschmidt, H. (2001). Speech perception deficit in dyslexic adults as measured by mismatch negativity (MMN). International Journal of Psychophysiology, 40(1), 77–87.

Schweinberger, S. R., & Neumann, M. F. (2016). Repetition effects in human ERPs to faces. Cortex, 80, 141–153.

Sederberg, P. B., Kahana, M. J., Howard, M. W., Donner, E. J., & Madsen, J. R. (2003). Theta and gamma oscillations during encoding predict subsequent recall. Journal of Neuroscience, 23(34), 10809–10814.

Seidenberg, M. S., & McClelland, J. L. (1989). A distributed, developmental model of word recognition and naming. Psychological Review, 96(4), 523.

Sigurdardottir, H. M., Danielsdottir, H. B., Gudmundsdottir, M., Hjartarson, K. H., Thorarinsdottir, E. A., & Kristjánsson, Á. (2017). Problems with visual statistical learning in developmental dyslexia. Scientific Reports, 7(1), 606.

Sigurdardottir, H. M., Fridriksdottir, L. E., Gudjonsdottir, S., & Kristjánsson, Á. (2018). Specific problems in visual cognition of dyslexic readers: Face discrimination deficits predict dyslexia over and above discrimination of scrambled faces and novel objects. Cognition, 175, 157–168.

Snyder, A. C., & Foxe, J. J. (2010). Anticipatory attentional suppression of visual features indexed by oscillatory alpha-band power increases: A high-density electrical mapping study. Journal of Neuroscience, 30(11), 4024–4032.

Sperling, A. J., Lu, Z. L., Manis, F. R., & Seidenberg, M. S. (2005). Deficits in perceptual noise exclusion in developmental dyslexia. Nature Neuroscience, 8(7), 862.

Starr, M. S., & Rayner, K. (2001). Eye movements during reading: Some current controversies. Trends in Cognitive Sciences, 5(4), 156–163.

Stoodley, C. J., Hill, P. R., Stein, J. F., & Bishop, D. V. (2006). Auditory event-related potentials differ in dyslexics even when auditory psychophysical performance is normal. Brain Research, 1121(1), 190–199.

Stoodley, C. J., Ray, N. J., Jack, A., & Stein, J. F. (2008). Implicit learning in control, dyslexic, and garden-variety poor readers. Annals of the New York Academy of Sciences, 1145, 173–183.

Strauss, A., Wöstmann, M., & Obleser, J. (2014). Cortical alpha oscillations as a tool for auditory selective inhibition. Frontiers in Human Neuroscience, 8, 350.

Summerfield, C., & Egner, T. (2009). Expectation (and attention) in visual cognition. Trends in Cognitive Sciences, 13(9), 403–409.

Summerfield, C., Trittschuh, E. H., Monti, J. M., Mesulam, M. M., & Egner, T. (2008). Neural repetition suppression reflects fulfilled perceptual expectations. Nature Neuroscience, 11(9), 1004.

Summerfield, C., Wyart, V., Mareike Johnen, V., & De Gardelle, V. (2011). Human scalp electroencephalography reveals that repetition suppression varies with expectation. Frontiers in Human Neuroscience, 5, 67.

Todorovic, A., & de Lange, F. P. (2012). Repetition suppression and expectation suppression are dissociable in time in early auditory evoked fields. Journal of Neuroscience, 32(39), 13389–13395.

Todorovic, A., van Ede, F., Maris, E., & de Lange, F. P. (2011). Prior expectation mediates neural adaptation to repeated sounds in the auditory cortex: An MEG study. Journal of Neuroscience, 31(25), 9118–9123.

Torgesen, J.K., Wagner, R.K., & Rashotte, C.A. (1999). TOWRE: Test of word reading efficiency Austin, TX: Pro-Ed.

Van der Lubbe, R. H., de Kleine, E., & Rataj, K. (2019). Dyslexic individuals orient but do not sustain visual attention: Electrophysiological support from the lower and upper alpha bands. Neuropsychologia, 125, 30–41.

Vidyasagar, T. R., & Pammer, K. (2010). Dyslexia: A deficit in visuo-spatial attention, not in phonological processing. Trends in Cognitive Sciences, 14(2), 57–63.

Vidyasagar, T. R. (2019). Visual attention and neural oscillations in reading and dyslexia: Are they possible targets for remediation? Neuropsychologia, 130, 59–65.

Vizioli, L., Rousselet, G. A., & Caldara, R. (2010). Neural repetition suppression to identity is abolished by other-race faces. Proceedings of the National Academy of Sciences, 107(46), 20081–20086.

Vogels, R. (2016). Sources of adaptation of inferior temporal cortical responses. Cortex, 80, 185–195.

Wacongne, C., Changeux, J. P., & Dehaene, S. (2012). A neuronal model of predictive coding accounting for the mismatch negativity. Journal of Neuroscience, 32(11), 3665–3678.

Wagner, R. K., Torgesen, R. W., & Rashotte, C. A. (1999). Comprehensive Test of Phonological Processing (CTOPP). Austin, TX: Pro-Ed.

Wechsler, D. (1999). Wechsler Abbreviated Scale of Intelligence. San Antonio, TX: Psychological Corporation.

Wechsler, D. (2008). Wechsler Adult Intelligence Scale – Fourth Edition. San Antonio, TX: Psychological Corporation.

White, T. P., Jansen, M., Doege, K., Mullinger, K. J., Park, S. B., Liddle, E. B., … & Liddle, P. F. (2013). Theta power during encoding predicts subsequent memory performance and default mode network deactivation. Human Brain Mapping, 34(11), 2929–2943.

Wiggs, C. L., & Martin, A. (1998). Properties and mechanisms of perceptual priming. Current Opinion in Neurobiology, 8(2), 227–233.

Woodcock, R. W., McGrew, K. S., & Mather, N. (2001). Woodcock-Johnson III Tests of Achievement. Itasca, IL: Riverside.

Woodcock, R. W. (1998). Woodcock Reading Mastery Tests – Revised/Normative Update. Circle Pines, MN: American Guidance Service.

Yeatman, J. D., Dougherty, R. F., Rykhlevskaia, E., Sherbondy, A. J., Deutsch, G. K., Wandell, B. A., & Ben-Shachar, M. (2011). Anatomical properties of the arcuate fasciculus predict phonological and reading skills in children. Journal of Cognitive Neuroscience, 23(11), 3304–3317.

Zhang, F., Jaffe Dax, S., Wilson, R. C., & Emberson, L. L. (2019). Prediction in infants and adults: A pupillometry study. Developmental Science, 22(4), e12780.

Ziegler, J. C., Pech Georgel, C., George, F., & Lorenzi, C. (2009). Speechuperceptionuinunoise deficits in dyslexia. Developmental Science, 12(5), 732–745.

