## Supplementary Materials for "Electrophysiological correlates of perceptual prediction error are attenuated in dyslexia"

**Supplementary Table 1. Expectation effects, as defined in the Dyslexia group.**

| **Cluster** | **Time (ms)** | **Freq. (Hz)** | **Distribution** | **Polarity** | **ANOVA Results** | ***F*_1,38_** | ***p*** |  |
| --- | --- | --- | --- | --- | --- | --- | --- | --- |
| ***Faces*** |  |  |  |  |  |  |  |  |
| #1 | 150–850 | 8–9 | global | ExpChg > ExpRep | *Group*  *Expectation*  *Group × Expectation* | 2.35  38.90  1.38 | 0.1  < 0.0001  0.2 | *** |
| #2 | 800–900 | 13 | global | ExpChg > ExpRep | *Group*  *Expectation*  *Group × Expectation* | 0.21  38.42  0.86 | 0.6  < 0.0001  0.4 | *** |
| #3 | 950–1250 | 8–9 | global | ExpChg > ExpRep | *Group*  *Expectation*  *Group × Expectation* | 0.42  34.31  0.24 | 0.5  < 0.0001  0.6 | *** |
| ***Words*** |  |  |  |  |  |  |  |  |
| #1 | 200–350 | 11–12 | global | ExpChg > ExpRep | *Group*  *Expectation*  *Group × Expectation* | 0.02  31.44  0.45 | 0.9  < 0.0001  0.5 | *** |
| #2 | 600–800 | 8 | posterior | ExpChg > ExpRep | *Group*  *Expectation*  *Group × Expectation* | 1.40  13.63  5.37 | 0.2  0.0007  0.03 | ***  * |

Clusters are numbered chronologically. Cluster time ranges are given with respect to the onset of S1. ***Abbreviations*** as in Table 2 in the main text.

**Supplementary Table 2. Repetition effects, as defined in the Dyslexia group.**

| **Cluster** | **Time (ms)** | **Distribution** | **Polarity** | **ANOVA Results** | ***F*_1,38_** | ***p*** |  |
| --- | --- | --- | --- | --- | --- | --- | --- |
| ***Faces*** |  |  |  |  |  |  |  |
| n/a | n/a | n/a | n/a | *n/a* |  |  |  |
| ***Words*** |  |  |  |  |  |  |  |
| #1 | 365–367 | posterior | Rep > Chg | *Group*  *Repetition*  *Group × Repetition* | 0.74  12.79  1.33 | 0.4  0.001  0.3 | ** |
| #2 | 379–520 | global | Rep > Chg | *Group*  *Repetition*  *Group × Repetition* | 0.09  90.35  0.08 | 0.8  < 0.0001  0.8 | *** |
| #3 | 527–539 | posterior | Rep > Chg | *Group*  *Repetition*  *Group × Repetition* | 0.32  9.71  4.33 | 0.6  0.003  0.04 | **  * |
| #4 | 553–561 | posterior | Rep > Chg | *Group*  *Repetition*  *Group × Repetition* | 0.48  9.38  1.42 | 0.5  0.004  0.2 | **** |

Clusters are numbered chronologically. Cluster time ranges are given with respect to the onset of S2. ***Abbreviations*** as in Table 4 in the main text.

**Supplementary Table 3. Prediction error, as defined in the Dyslexia group.**

| **Cluster** | **Time (ms)** | **Distribution** | **Polarity** | **ANOVA Results** | **F_1,38_** | ***p*** |  |
| --- | --- | --- | --- | --- | --- | --- | --- |
| ***Faces*** |  |  |  |  |  |  |  |
| n/a | n/a | n/a | n/a | *n/a* |  |  |  |
| ***Words*** |  |  |  |  |  |  |  |
| #1 | 356–357 | central | Rep > Chg | *Group*  *Repetition*  *Expectation*  *Group × Repetition*  *Group × Expectation*  *Repetition × Expectation*  *Group × Repetition × Expectation* | 0.01  34.93  10.29  0.02  2.87  0.80  0.27 | 0.9  < 0.0001  0.003  0.9  0.1  0.4  0.6 | ***  ** |
| #2 | 383–414 | global | Rep > Chg | *Group*  *Repetition*  *Expectation*  *Group × Repetition*  *Group × Expectation*  *Repetition × Expectation*  *Group × Repetition × Expectation* | 0.01  89.42  9.80  0.40  1.70  4.57  1.34 | 0.9  < 0.0001  0.003  0.5  0.2  0.04  0.3 | **  **  * |
| #3 | 420–432 | central | Rep > Chg | *Group*  *Repetition*  *Expectation*  *Group × Repetition*  *Group × Expectation*  *Repetition × Expectation*  *Group × Repetition × Expectation* | 0.02  66.59  3.78  0.00  0.62  7.25  0.28 | 0.9  < 0.0001  0.06  1.00  0.4  0.01  0.6 | ***  * |

Clusters are numbered chronologically. Cluster time ranges are given with respect to the onset of S2. ***Abbreviations*** as in Table 5 in the main text.
